# Profiling of human neural crest chemoattractant activity as replacement of fetal bovine serum for in vitro chemotaxis assays

**DOI:** 10.1101/2021.07.19.452897

**Authors:** Xenia Dolde, Christiaan Karreman, Marianne Wiechers, Stefan Schildknecht, Marcel Leist

## Abstract

Fetal bovine serum (FBS) is the only known stimulus for migration of human neural crest cells (NCCs). Non-animal chemoattractants are desirable for the optimization of chemotaxis assays to be incorporated in a test battery for reproductive and developmental toxicity. We confirmed here in an optimized transwell assay that FBS triggers directed migration along a concentration gradient. The responsible factor was found to be a protein in the 30-100 kDa size range. In a targeted approach, we tested a large panel of serum constituents known to be chemotactic for NCCs in animal models (e.g. VEGF, PDGF, FGF, SDF-1/CXCL12, ephrins, endothelin, Wnt, BMPs). None of the corresponding human proteins showed any effect in our chemotaxis assays based on human NCCs. We then examined, whether human cells would produce any factor able to trigger NCC migration in a broad screening approach. We found that HepG2 hepatoma cells produced chemotaxis-triggering activity (CTA). Using chromatographic methods and by employing the NCC chemotaxis test as bioassay, the responsible protein was enriched by up to 5000-fold. We also explored human serum and platelets as direct source, independent of any cell culture manipulations. A CTA was enriched from platelet lysates several thousand-fold. Its temperature and protease-sensitivity suggested also a protein component. The capacity of this factor to trigger chemotaxis was confirmed by single-cell video-tracking analysis of migrating NCCs. The human CTA characterized here may be employed in the future for the setup of assays testing for the disturbance of directed NCC migration by toxicants.

## Introduction

The directed migration of neural crest cells (NCCs) over large distances is essential for normal vertebrate development. Genetic defects interfering with this process can lead to a broad panel of malformations and disease syndromes, such as Hirschsprung’s disease, Treacher Collins syndrome or Waardenburg syndrome (Vega-Lopez et al., 2018; Serrano et al., 2019). Chemicals that interfere with NCC migration often lead to craniofacial defects in the developing fetus (Zhang et al., 2017). This is well documented for ethanol or pesticides like triadimefon (Menegola et al., 2005). Disturbed retinoic acid (RA) levels are an important cause of impaired NCC migration and differentiation. Under such conditions, craniofacial defects are observed in both, animals and humans (Williams and Bohnsack, 2019).

NCCs are multipotent cells generated at the lateral edges of the neural plate. During early fetal development, NCCs migrate long distances to their target sites such as the skin, the skull and the intestine. They differentiate into a large variety of cell types, including neurons, melanocytes and chondrocytes (Le Douarin, 2004). NCCs are grouped into subpopulations according to their position within the anteroposterior axis of the embryo. Cranial NCCs build mainly structures of the head (Prasad et al., 2019), cardiac NCCs contribute to the smooth muscle of the great vessel/aorta (Sieber-Blum, 2004), and trunk NCCs give rise to sensory neurons, the sympathoadrenal system and pigment cells (Giovannone et al., 2015; Huang et al., 2016).

Cell migration is a complex process, involving several biological functions, e.g. adhesion of cells to the extracellular matrix (ECM), detachment from the substrate, and remodeling of the cytoskeleton. During the migration process protrusions are extended at the leading edge, whereas the trailing edge is contracted and cell material is moved to the front pole of the cell (Conway and Jacquemet, 2019; Ridley et al., 2003). The process by which factors promote increased cell motility is called chemokinesis, whereas chemotaxis is defined as the guided movement of cells along a gradient of bound molecules, soluble factors or mechanical stimuli (Shellard and Mayor, 2016). To perform chemotaxis, cells need to have increased motility, but also display properties such as directional sensing and maintenance of polarity (Kay et al., 2008). Polarized cells are defined by a front that has localized actin polymerization and a rear that is able to contract (Kay et al., 2008). Directional sensing is the property of cells to compare receptor occupancy over their surface, and to determine where the concentration is the highest (Kay et al., 2008). In the presence of a chemoattractant gradient, the cells sense the gradient, align their polarity with it and finally migrate along the gradient (Wu, 2005).

Migration of NCCs is initiated by a process called epithelial-to-mesenchymal transition (EMT), which goes along with several motility increasing changes that affect cell polarity and adhesive properties (Nieto et al., 2016). Chemotaxis has been observed for individual NCCs, but also for groups of cells moving in a co-ordinated manner like e.g. wild geese (Capuana et al., 2020). Collective migration allows a cluster of NCCs to migrate faster and to follow a weak chemoattractant gradient, to which a single cell would be insensitive (Merchant and Feng, 2020; Mayor and Etienne-Manneville, 2016). A cluster of migrating cells is defined by leader and follower cells, which differ in their gene expression (Capuana et al., 2020). The transcriptome patterns controlling such behaviour are dependent on local environment, and experimental systems (McLennan et al., 2015), and it is likely that chemotactically active NCCs differ from cells not following a gradient.

Various factors have been proposed so far as NCC chemoattractants e.g. vascular endothelium growth factor (VEGF) for chicken cranial NCCs (McLennan et al., 2010) and platelet derived growth factor (PDGF) in cranial NCCs of the zebrafish (Eberhart et al., 2008), fibroblast growth factors (FGF) in the cranial, cardiac and trunk regions of mice (Kubota and Ito, 2000), and stromal cell derived factor 1 (SDF-1/CXCL12) in the cranial and trunk regions of chicken (Kasemeier-Kulesa et al., 2010). Current knowledge on NCC development has mainly been obtained from animal models. The most common *in vivo* or *in vitro* experiments to investigate NCC migration and chemotaxis were performed using *Xenopus laevis*, mouse, rat and chick NCCs (Bahm et al., 2017; Theveneau et al., 2010; McLennan et al., 2010; Kubota and Ito, 2000). Thus, most of the above mentioned chemoattractants have not been confirmed for human NCCs. Using human NCCs differentiated from human pluripotent stem cells (hPSCs) is slowly becoming attractive in the field and several differentiation protocols are available (Hackland et al., 2017; Hackland et al., 2019; Tchieu et al., 2017; Zimmer et al., 2012; Chambers et al., 2016). These *in vitro* differentiation protocols gave new insights into the molecular mechanisms of human NCC development. The use of induced pluripotent stem cells (iPSCs) from patients, has enabled disease modeling of neurocristopathies (Srinivasan and Toh, 2019; Workman et al., 2017; Lee et al., 2009; Zeltner et al., 2016). Moreover, experimental models based on human NCCs have helped to identify chemicals that inhibit migration (Zimmer et al., 2012; Nyffeler et al., 2017a). Unfortunately, data on consistent, concentration-dependent chemotaxis stimuli are still lacking. Such a stimulus would be required to test whether chemicals can specifically impair directed migration.

At the moment, there are thousands of untested chemicals used in commerce and an assessment of all their potentially-harmful properties in complex animal models is not feasible. Due to potential species differences, human cell-based high-throughput screening (HTS) methods are required (Hartung, 2009; Hartung and Leist, 2008; Collins et al., 2008). Such new approach methods (NAMs) should allow for the cheap and fast testing of many chemicals (Crofton et al., 2012; Aschner et al., 2017; Bal-Price et al., 2018; Krebs et al., 2020; Smirnova et al., 2014). A NCC chemotaxis assay could be incorporated in a NAM test battery (Zimmer et al., 2014) and used in the context of next generation risk assessment (NGRA) (Baltazar et al., 2020; Vinken et al., 2021). Based on this, animal-free risk assessment for the safety of compounds may be performed (Moné et al., 2020).

Indeed, several *in vitro* assays to investigate NCC migration have been established during the past ten years based on human NCCs differentiated from hPSCs (Lee et al., 2010; Zimmer et al., 2012). In the original wound healing assay, a scratch was introduced in an NCC monolayer to create a cell-free area. Many toxicants interfering with the movement of cells into the gap have been identified, and the circular migration inhibition of neural crest cell (cMINC) assay is an improved version (robustness and throughput) of the original scratch assay. As in the original wound healing assay, the NCCs migrate in a random manner into the cell-free zone (Nyffeler et al., 2017b; Nyffeler et al., 2017a). Known NCC toxicants were confirmed (including valproic acid (VPA), methylmercury chloride, As2O3, CdCl2 and polychlorinated biphenyls (PCBs)) and several unknown hazardous chemicals have been identified (Dreser et al., 2015; Zimmer et al., 2012; Zimmer et al., 2014; Nyffeler et al., 2017b; Nyffeler et al., 2018).

The above mentioned assays model NCC migration, but they are not able to assess directed cell migration. Chemotaxis assays require a stable gradient of a chemoattractant, which can be sensed by the cells (Shellard and Mayor, 2016). To construct such a gradient there is an urgent need for a human NCC chemoattractant. To our knowledge, bovine serum is the only known stimulus of motility, described in the literature. Based on its animal origin and its poorly standardized composition, it is not the ideal basis for an assay set-up.

The aim of this study was therefore to identify better defined chemoattractants to study directed NCC migration. The study set out to verify that FBS indeed triggers chemotaxis, and not just chemokinesis. Further approaches were used to demonstrate that a protein factor is responsible for the chemotactic activity of FBS. Based on this knowledge, human cell lines were screened for their capacity to secrete such a factor and HepG2 cells were found as suitable source. As alternative, pure human starting material, platelet lysates were considered. They were found to contain a potent NCC chemoattractant, which was highly enriched in the course of the study.

## Materials and Methods

### Neural crest cell differentiation

NCCs were differentiated from the induced pluripotent stem cell (iPSC) line IMR90_clone_#4 (WiCell, Madison, Wisconsin, USA) following the modified protocol of Mica et al. (2013). IPSCs were maintained on human Laminin-521 (BioLamina, Sundbyger, Sweden) coating in essential 8 (E8) medium (DMEM/F12 supplemented with 15 mM Hepes (Gibco/Fisher Scientific, Hampton, New Hampshire, USA), 16 mg/ml, L-ascorbic-acid, 0.7 mg/ml sodium selenite, 20 µg/ml insulin, 10 µg/ml holo-transferrin (all from Sigma, Steinheim, Germany), 100 ng/ml bFGF (Thermo Fisher Scientific, Waltham, Massachusetts, US), 1.74 ng/ml TGFb (R&D Systems, Minneapolis, Minnesota, USA). For differentiation into NCCs, iPSCs were plated on Matrigel^TM^ (Corning, Glendale, Arizona, USA) coated 6-well plates at a density of 100,000 cells/cm^2^ in E8 medium containing 10 μM ROCK-inhibitor (Y-27632 (Tocris, Bristol, UK)). After one day cells reached a confluency of 70-80% and differentiation was initiated (day 0) by a medium change to KSR medium (Knock out DMEM, 15% knock out serum replacement, 1% GlutaMax, 1% MEM NEAA solution, 50 µM 2-mercaptoethanol; (all from Gibco/Fisher Scientific, Hampton, New Hampshire, USA)) supplemented with 20 ng/ml Noggin (R&D Systems, Minneapolis, Minnesota USA) and 10 µM SB431542 (Tocris, Bristol, UK). From day 2 on cells were treated with 3 µM CHIR 99021 (Axon Medchem, Reston, Virginia, USA). Noggin and SB431542 were withdrawn at day 3 and 4, respectively. Beginning at day 4, the KSR medium was gradually replaced by 25% increments of N2-S medium (DMEM/F12, 1% GlutaMax (both from Gibco/Fisher Scientific, Hampton, New Hampshire, USA), 1.55 mg/ml glucose, 0.1 mg/ml apotransferin, 25 µg/ml insulin, 20 nM progesterone, 100 μM putrescine, 30 nM selenium (all from Sigma, Steinheim, Germany)). Cells were collected at day 11, resuspended in N2-S medium supplemented with 20 ng/ml EGF and 20 ng/ml FGF2 (both from R&D Systems, Minneapolis, M USA) and seeded as droplets (10 µl) on poly-L-ornithine (PLO)/Laminin/Fibronectin (all from Sigma, Steinheim, Germany) coated 10 cm dishes. Cells were expanded by weekly splitting. From now on seeding as droplets was not necessary and medium was changed every second day. After 35-39 days, cells were cryopreserved at a concentration of 4*10^6^ cells/ml in 90% N2-S medium and 10% dimethyl sulfoxide (DMSO) (Merck Millipore, Burlington, Massachusetts, USA) until further use.

### Migration assay (cMINC)

The circular migration inhibition of neural crest cell (MINC) assay was performed as described earlier (Nyffeler et al., 2017b). Briefly, silicone stoppers (Platypus Technologies, Madison, WI, USA) were placed centrally into each experimental well of a 96-well polystyrene plate (Corning, Glendale, Arizona, USA) coated with 1 µg/ml fibronectin and 1 µg/ml laminin (both from Sigma, Steinheim, Germany). Cells were seeded around the stoppers at a density of 95,000 cells/cm^2^. The following day, stoppers were removed to allow cells to migrate into the cell-free central area and medium was refreshed. To test the effect of toxicants on NCC motility, 5x concentrated toxicant solution was added to the medium 24 h after stopper removal. After another 24 h, cell viability and migration endpoints were monitored. For this, cells were stained with HOECHST-33342 and calcein-AM (both from Sigma, Steinheim, Germany) and image acquisition was performed using a Cellomics ArrayScan VTI imaging microscope (Thermo Fisher, Pittsburgh, Pennsylvania, USA). HOECHST-33342 and calcein double positive cells were defined as viable cells and determined by an automated algorithm described earlier (Stiegler et al., 2011; Krug et al., 2013). For quantification of migration, a free software tool (http://invitrotox.uni-konstanz.de/RA/) was used as described in Nyffeler et al. (2017b) to calculate the original stopper position and determine the number of HOECHST-33342 and calcein double positive cells within the migration area. Viability and migration were normalized to untreated or solvent control (0.1% DMSO).

### Neural crest membrane translocation (NC-MT) assay

For the NC-MT assay, Transwell® 24 well permeable supports (pore size 8 µm, polycarbonate membrane, Corning, Glendale, Arizona, USA, catalog no. 3422) were used. NCCs were seeded at a density of 50’000 cells per insert (150,000 cells/cm^2^, 100 µl) in N2-S medium supplemented with 20 ng/ml EGF and 20 ng/ml FGF2 (both from R&D Systems, Minneapolis, Minnesota, USA) into the upper chamber. Test compounds were added in the stated concentration to the lower chamber (650 µl). The cells were allowed to migrate for 6 h at 37 °C and 5% CO2. After incubation, medium was aspirated from inserts and reservoirs and the upper side of each insert was gently swabbed, using cotton-swabs, to remove cells that had not migrated through the membrane. Reservoirs and inserts were washed once with phosphate buffered saline (PBS) and afterwards the migrated cells on the membrane were fixed with 3.7% formaldehyde (v/v in H2O) and stained with crystal violet for 15 min. Then, the inserts were thoroughly rinsed with water and dried for at least 24 h. Five pictures per condition were taken with an Axio Observer Z1 microscope (Zeiss, Oberkochen, Germany) to evaluate the number of migrated cells. The number of migrated cells was normalized to that of cells stimulated with FBS.

For the NC-MT-HTS assay, the Transwell® high troughput screening system (pore size 8 µm, polyester membrane, Corning, Glendale, Arizona, USA, catalog no. 3384) was used. It consists of 96 wells of permeable inserts connected by a rigid tray and a 96 well receiver plate. Cells were seeded at a density of 25’000 cells per insert (175’000 cells/cm^2^, 50 µl) into the upper chamber. Test compounds were added in the stated concentration to the lower chamber (150 µl). The cells were allowed to migrate for 6 h at 37 °C and 5% CO2. After incubation, medium was aspirated from inserts and reservoirs and both were washed once with PBS. Reservoirs were filled with 150 µl EDTA solution containing calcein-AM (Sigma, Steinheim, Germany) and incubated for 30 min at 37 °C and 5% CO2. Plates were then centrifuged for 4 min at 350 x g to remove migrated cells from the membrane. The tray with 96 wells of permeable inserts was removed and the receiver plate containing the migrated cells was placed into a spectrophotometer (TECAN, Männedorf, Switzerland). Calcein-AM staining was detected at an emission length of 520 nm. After subtraction of the blank values, the number of migrated cells was normalized to that of cells stimulated with FBS.

### Determination of chemotactic behaviour by cell tracking

The µ-slide chemotaxis assay (Ibidi, Martinsried, Germany) allows the establishment of a stable gradient and the observation of cells within this gradient via time-lapse imaging. The two reservoirs, filled either with a chemoattractant or with medium, are connected by a gap. The gap was coated with 1 µg/ml fibronectin (Sigma, Steinheim, Germany) one day before seeding of the cells. The gap was filled with 6 µl of cell suspension with a concentration of 3x10^6^ cells/ml (= 18,000 cells). The cells were allowed to attach for 3 h at 37 °C and 5% CO2. Afterwards, the reservoirs were filled with pure medium on the one side and medium containing chemoattractant on the other side. Right afterwards, the µ-slide chemotaxis slide was mounted on the stage of an Axio Observer Z1 microscope (Zeiss, Oberkochen, Germany) equipped with an Axiocam MRm camera and an incubation chamber (37 °C, 5% CO2). Phase contrast images were taken for 24 h every 10 minutes using a 5x objective. Images were exported as JPEG files and cell tracking was performed using the ‘Manual Tracking’ plugin from ImageJ (Schneider et al., 2012). For each biological replicate, 20 cells were tracked per condition. The resulting cell coordinates were transferred to the ‘Chemotaxis and Migration Tool V2.0’ (Ibidi, Martinsried, Germany) to determine cell translocation, accumulated distance and cell speed and to generate “rose plots” of the tracked cells.

### Co-culture and conditioned medium preparation

HepG2 hepatoma cells (ATCC, HB-8065), MDA-MB-231 breast carcinoma cells (ATCC, HTB- 26), HeLa cervical cancer cells (ATCC, CCL-2), HEK-239 human embryonic kidney cells (ATCC, CRL-1573) and SH-SY5Y neuroblastoma cells (ATCC, CRL-2266) were cultured in DMEM + GlutMax^TM^ (Gibco/Fisher Scientific, Hampton, New Hampshire, USA) supplemented with 10% foetal bovine serum (FBS) (PAA Laboratories, Pasching, Austria) and 1% pen/strep (Gibco/Fisher Scientific, Hampton, New Hampshire, USA) at 37 °C and 5% CO2. Cells were passaged every other day.

For conditioned medium (CM) preparation, the cells were seeded in DMEM + GlutMax^TM^ (Gibco/Fisher Scientific, Hampton, New Hampshire, USA) supplemented with 10% FBS (PAA Laboratories, Pasching, Austria) and 1% pen/strep (Gibco/Fisher Scientific, Hampton, New Hampshire, USA) and grown until they reached confluency. Then, medium was aspirated, the cells were washed once with phosphate buffered saline (PBS) and then fresh DMEM + GlutMax^TM^ (Gibco/Fisher Scientific, Hampton, New Hampshire, USA) without FBS and pen/strep was added. Cells were incubated for 24 h. To remove any residues of FBS, medium was again aspirated, the cells were washed once with PBS, and then fresh DMEM + GlutMax^TM^ (Gibco/Fisher Scientific, Hampton, New Hampshire, USA) was added. After incubation for another 24 h, the medium supernatant was harvested and centrifuged at 314 x g for 4 min, to remove cell debris. This conditioned medium (CM) was either used in the transwell assay or further processed by acetone precipitation (pellet stored at -20 °C).

For co-culture experiments, cells were seeded at a concentration of 150’000 cells/cm^2^ in 24-well plates in DMEM + GlutMax^TM^ (Gibco/Fisher Scientific, Hampton, New Hampshire, USA) supplemented with 10% foetal bovine serum (FBS) (PAA Laboratories, Pasching, Austria) and 1% pen/strep (Gibco/Fisher Scientific, Hampton, New Hampshire, USA). Cells were incubated for 24 h. Afterwards, cells were washed once with phosphate buffered saline (PBS) and then fresh DMEM + GlutMax^TM^ (Gibco/Fisher Scientific, Hampton, New Hampshire, USA) was added. The steps were analogue to the conditioned medium preparation. After 48 h of starvation, the transwell inserts with seeded NCCs were placed into the wells of the 24-well plate containing starved cells.

### Human serum and platelet lysate preparation

Approach 1 (human serum): Whole Blood was obtained from healthy adult volunteers and collected into Monovettes (7.5 ml, K3 EDTA, Sarstedt, Nümbrecht, Germany). Procedures were approved by the institutional review board (IRB) of the University of Konstanz. Before blood collection, the monovettes were washed twice with MilliQ water to remove EDTA and enable clotting. For serum preparation, whole blood was allowed to clot by leaving it undisturbed at RT for 30 min. The clot was removed by centrifugation at 1500 x g for 10 min at 4 °C. Supernatants were transferred immediately into fresh tubes, aliquoted and stored at -80 °C.

Approach 2 (huPL preparation): For platelet lysate preparation, whole blood was centrifuged at 150 g for 20 min at RT to collect platelet rich plasma. Only the upper half of the platelet rich plasma was transferred into a new plastic tube and buffer A (10 mM sodium citrate, 150 mM NaCl, 1 mM EDTA, 1% dextrose, pH 7.4) containing 1 µM prostaglandin (PGI2, Iloprost, Sigma, Steinheim, Germany) was added at 1:1 ratio. The mixture was centrifuged at 350 x g for 15 min at RT. The platelet pellet was washed once in buffer B (140 mM NaCl, 6 mM KCl, 2 mM Mg2SO4, 2 mM NaHPO4, 6 mM HEPES, pH 7.4) to remove plasma residues. The pellet was resuspended in an appropriate volume of buffer B and the platelets were lysed by three freeze-thaw cycles at -20 °C and 37 °C. Finally, the platelet lysate was centrifuged at 2000 x g for 15 min at RT to remove platelet debris. Supernatant was aliquoted and stored at -20 °C.

Approach 3: Commercial huPLs (CRUX RUFA research (Trinova Biochem, Gießen, Germany), ELAREM™ PRIME (PL BioScience, Aachen, Germany), Stemulate™ (COOK Regentec, Indianapolis, Indiana, USA)) were obtained, aliquoted and stored at -20 °C. To prevent coagulation of the cell culture medium, heparin (PL-HEP-0005) was added at a final concentration of 2 U/ml. No heparin was required when using Stemulate™ (COOK Regentec, Indianapolis, Indiana, USA).

### Acetone precipitation

FBS, huPL or HepG2 CM were mixed with 30% precooled acetone (VWR Chemicals, Darmstadt, Germany) and incubated overnight at -20 °C. Samples were then centrifuged at 7000 x g for 30 min. Afterwards, the supernatant was transferred into a new 250 ml centrifuge tube (Corning, Glendale, Arizona, USA) and the pellet was discarded. Then 10% precooled acetone (VWR Chemicals, Darmstadt, Germany) was added and again incubated overnight at -20 °C. The sample was centrifuged at 7000 x g for 30 min. This time the supernatant was discarded and the remaining pellet dried at room temperature until acetone remains were evaporated. The pellet was stored at - 20 °C for further usage.

### Protein purification

Fast protein liquid chromatography (FPLC) was used to purify complex protein mixtures. The procedure is taking advantage of the fact, that different proteins have different affinities to the resin of the purification columns. FPLC was performed using an ÄKTAprime plus (GE Healthcare, München, Germany) system equipped with a UV detection system. Protein separation was carried out with different ion exchange columns: A cation exchange column (HiScreen Capto SP ImpRes, GE Healthcare, München, Germany), an anion exchange column (HiTrap Q HP 1 ml, GE Healthcare) and another anion exchange column (HiTrap Q FF 5 ml, GE Healthcare, München, Germany). The start buffer contained 10 mM Tris-HCl (pH 7.4) and the elution buffers contained additionally 2 M MgCl2 for the cation exchange column or 2 M NaCl for the anion exchange columns. All buffers were sterile-filtered before usage. The chromatographic separation was performed using a linear gradient of the elution buffer, starting from 0 up to 50%, followed by a step up to 100% (**Figure S5**). The protein sample was sterile-filtered before loading it to the column. FPLC separation was performed at room temperature (RT) and fractions of 1 ml were separated with a fraction collector. Within the stepwise purification process, the collected fractions were tested for bioactivity in the NC-MT assay. For this, fractions were diluted 1 + 4 with medium and the migration increasing activity was tested. For further purification steps, the active fractions were combined and desalted with a desalting column (HiPrep 26/10 Desalting, GE Healthcare, München, Germany), using MilliQ water as elution buffer. Afterwards, the desalted sample was loaded on the second ion exchange column for further protein separation. The individual fractions were tested in the NC-MT assay for their migration increasing activity.

### Protein separation and detection

For polyacrylamide gel electrophoresis (PAGE), samples were lysed in 1x Laemmli buffer and boiled for 5 minutes at 95 °C. Thirty-five micrograms of total protein were loaded on 10% PAA gels. The gel was then stained with coomassie blue dye (InstantBlue, VWR Chemicals, Darmstadt, Germany) for 20 min or overnight and afterwards washed twice with desalted water. Silver stainings were performed according to the instruction manual of the Pierce™ Silver Stain for Mass Spectrometry kit (Pierce/Thermo Fisher Scientific, Rockford, Illinois, USA, catalog no. 24600).

For western blotting, samples were lysed in 1x Laemmli buffer and boiled for 5 minutes at 95 °C. Thirty-five micrograms of total protein were loaded on 10% PAA gels. Afterwards, proteins were transferred onto nitrocellulose membranes (Amersham, Buckinghamshire, UK) using the Invitrogen iBlot 2 system. Membranes were blocked with 5% milk powder (w/v) in 0.1% TBS-Tween (v/v) for at least 1 h. Fibronectin antibody ab45688 (Abcam, Cambridge, UK), alpha 1 fetoprotein (AFP) antibody ab133617 (Abcam, Cambridge, UK) and glyceraldehyde 3-phosphate dehydrogenase (GAPDH) antibody ZG003 (Thermo Fisher Scientific, Waltham, Massachusetts, US) were incubated at 4 °C overnight. After washing steps with 0.1% TBS-Tween (v/v), secondary antibody (peroxidase-conjugated AffiniPure goat anti-mouse IgG (Jackson Immunoresearch, Cambridge, UK) or horseradish peroxidase-conjugated donkey anti-rabbit IgG (GE Healthcare, München, Germany)) was incubated for 1 h at RT. For visualisation, ECL western blotting substrate (Pierce/Thermo Fisher Scientific, Rockford, IL, USA) was used.

For protein quantification, the Quick Start^TM^ Bradford protein assay Kit (Bio Rad, München, Germany) was used and the assay was performed according to the instruction manual of the manufacturer. BSA was used as protein standard with concentrations from 2 mg/ml down to 1.25 µg/ml. The assay was performed in a 96-well plate and BSA standard dilutions and sample dilutions were added in triplicates to the wells, respectively. After incubation for 5 min at room temperature, the absorbance was measured at 595 nm with a spectrophotometer (TECAN, Männedorf, Switzerland).

### Mass spectrometry (MS) for protein identification

The active fractions from the NC-MT assay were separated in a 10% SDS gel and stained with coomassie blue (InstantBlue, VWR Chemicals, Darmstadt, Germany) or silver stain kit for mass spectrometry (Pierce/Thermo Fisher Scientific, Rockford, Illinois, USA, catalog no. 24600). The protein bands of interest were cut out and all further steps were performed at the Proteomics Centre of the University of Konstanz.

For sample preparation, all samples were reduced with DTT (30min, 56°C) and alkylated with chloroacetamide (60min, RT). Digestions were performed using Trypsin (16h, 30°C).

All digests were analysed on a QExactive HF mass spectrometer (Thermo Fisher Scientific, Bremen, Germany) interfaced with an Easy-nLC 1200 nanoflow liquid chromatography system (Thermo Fisher Scientific, Bremen, Germany). The peptide digests were reconstituted in 0.1% formic acid and loaded onto the analytical column (75 μm × 15 cm). Peptides were resolved at a flow rate of 300 nL/min using a linear gradient of 6−45% solvent B (0.1% formic acid in 80% acetonitrile) over 45 min. Data-dependent acquisition with full scans in a 350− 1500 m/z range was carried out at a mass resolution of 120,000. The 15 most intense precursor ions were selected for fragmentation. Peptides with charge states 2−7 were selected, and dynamic exclusion was set to 30 sec. Precursor ions were fragmented using higher-energy collision dissociation (HCD) set to 28%. For data evaluation, the raw data were searched against a suitable database using Proteome Discoverer 1.4 (Thermo Scientific).

### Heat inactivation and pepsin digestion

FBS, huPL and HepG2 CM samples were heat treated under different conditions. Therefore, the samples were incubated in heat blocks with temperatures of 60 °C and 70 °C for 30 min and in heat blocks with temperatures of 80 °C and 90 °C for 15 min.

For pepsin digestion FBS, huPL and HepG2 CM samples were mixed with pepsin solution, resulting in a final concentration of 0.5% pepsin (w/v). The pH was adjusted to pH=2 with a 1 M HCl solution and controlled with pH strips. The samples were incubated at 37 °C in a waterbath. After 1 h of incubation, the pH of the sample was adjusted to pH=7 by adding 1 M NaOH solution to stop the pepsin reaction. The control sample was run through the same acidification-incubation procedure, but without pepsin. Control samples were prepared for each condition to exclude the influence of pH change on the sample activity.

### Stability control tests

Collected fractions after ion exchange chromatography were tested directly in the NC-MT assay for their migration increasing activity. Afterwards, the remaining fractions were stored for 24 h under different conditions: samples were left at 4 °C or frozen at -80 °C, samples were mixed with 1 mM protease inhibitor (cOmplete tablets, Roche, Basel, Switzerland) and stored at 4 °C, samples were shock frozen in liquid nitrogen and stored at -80 °C, samples were mixed with 0.5 % BSA and stored at 4 °C and -80 °C, and samples were mixed with 20% glycerol and stored at -80 °C. After 24 h of incubation, the samples were retested in the NC-MT assay for their migration increasing activity.

### Fractionation of protein according to their molecular weight

Centrifugal filter devices (Amicon®- Ultra-0.5, Merck Millipore, Burlington, Massachusetts, USA) in five different “cut-off” sizes (3 K, 10 K, 30 K, 50 K, 100 K; K=1000 Da) were filled with 500 µl sample and centrifuged for 15 min at 14,000 x g at RT. The collection tube (filtrate) contained proteins smaller than the molecular cut-off. Proteins larger than the cut-off remained after centrifugation in the filter device (supernatant). To recover the sample in the filter device, the device was placed upside down in an empty tube and centrifuged for two minutes at 1,000 x g at RT.

### Data handling and statistics

If not stated otherwise, values are expressed as means of at least three different experiments (i.e. using three different cell preparations), with at least three technical replicates per cell preparation. Statistical differences were tested by ANOVA with post hoc tests as appropriate, using GraphPad Prism 7.0 (Graphpad Software, La Jolla, USA, www.graphpad.com).

## Results and Discussion

### Establishment of a chemotaxis assay based on human NCCs

FBS has been shown earlier to accelerate the mobility of NCCs (Nyffeler et al., 2017b), and it was therefore used here as a promising first candidate to establish a chemoattractant gradient for a chemotaxis assay. As assay principle, we used a modified Boyden chamber approach (Boyden, 1962; Mastyugin et al., 2004). Moreover, this setup has earlier been proven useful for studying inhibition of the movement of NCCs by test chemicals (Nyffeler et al., 2017a; Nyffeler et al., 2018). As cell source, we used NCCs differentiated from pluripotent stem cells. Such cells have been characterized by comprehensive transcriptome analysis, and they have been used successfully as test system for *in vitro* assays (Nyffeler et al., 2017a; Pallocca et al., 2017; Zimmer et al., 2014; Klose et al., 2021; Lee et al., 2010).

The main advantage of our assay is that cells (NCCs) are cultured on top of a porous membrane in the upper compartment of a two chamber system, and that migration of the cells through the membrane into the lower chamber can be easily quantified. For this reason, we termed our test: “neural crest-membrane translocation” (NC-MT) assay. In this system, a chemoattractant gradient can be established across the membrane by adding different concentrations of chemoattractant into the upper and lower chamber (**Figure 1A**). It was shown that the cells stay on the (lower surface of the) membrane, once they have migrated. Therefore, quantification of migration was very straightforward: cells were stained and counted at the end of the migration period (6 h) (**Figure 1B**). In order to verify that FBS triggers true chemotaxis, various experimental conditions were compared. Cells migrated only when a gradient across the membrane was established and they sensed a higher FBS concentration in the lower compartment. Direct contact of cells to high concentrations of FBS in the upper compartment did not trigger migration across the membrane towards the lower compartment. We therefore conclude that the assay assesses genuine chemotaxis (**Figure 1C**). In this setup, many NCC chemoattractants known from animal cell studies were tested. None of them showed a chemotactic effect on human NCCs in the NC-MT assay (**Figure 1D**). Using the same assay, we found a chemotaxis-triggering activity (CTA) in human serum (huSerum), which was similar to the one in FBS. Thus, also human serum contains a NCC chemoattractant and can be used for assay setup (**Figure 1D**).

**Figure 1:**
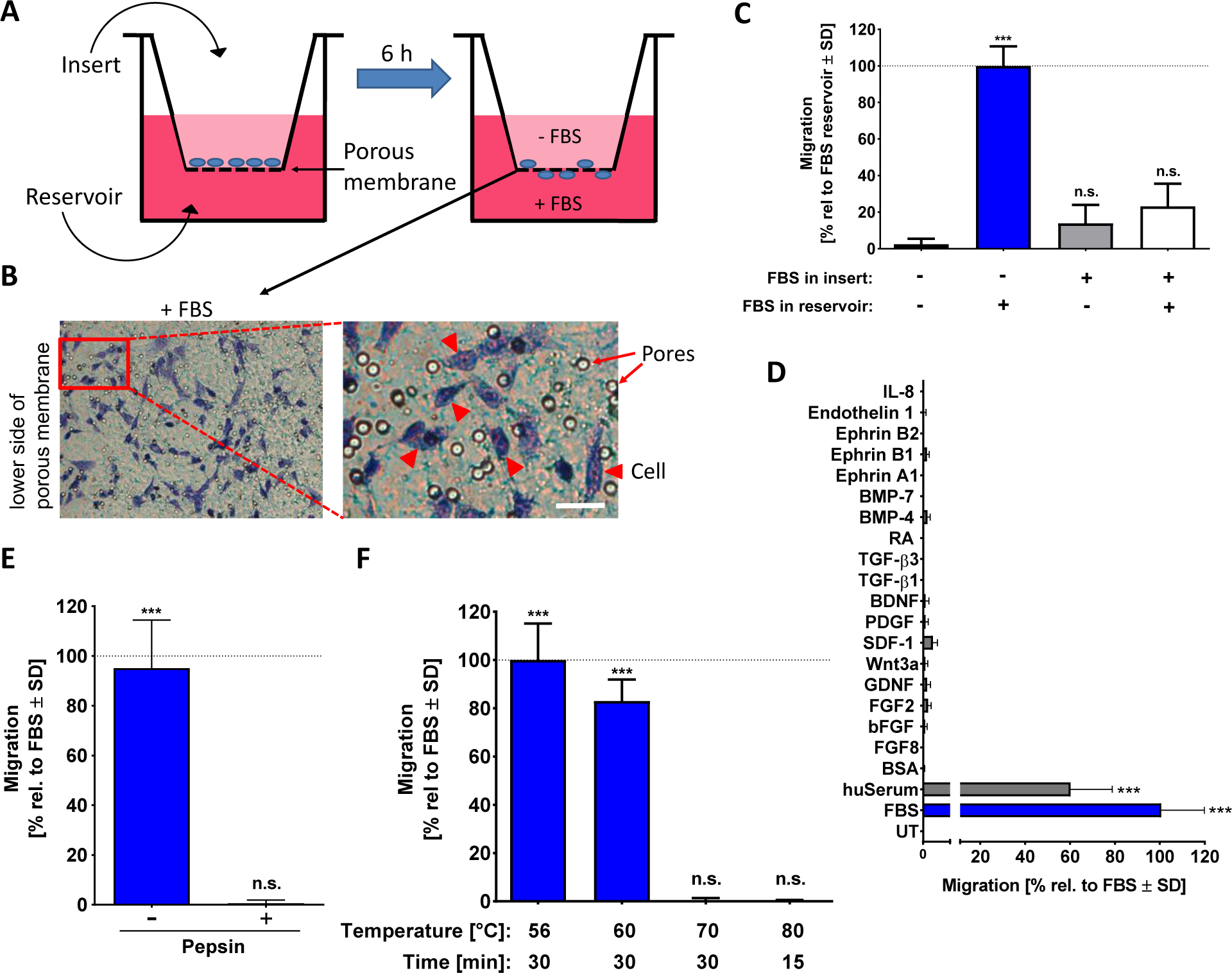
Characterization of the chemotaxis-promoting factor of bovine serum. (A) Graphical representation of the neural crest-membrane translocation (NC-MT) assay principle. Cells were plated into cell culture inserts equipped with a porous membrane at the bottom. This allows the addition of different amounts of potential chemoattractants to the reservoir and insert compartments. A chemoattractant gradient is thereby formed in the pores of the membrane. (B) FBS is shown as an example of a compound that triggers NCCs to migrate through the pores and to settle on the lower surface of the membrane. Example images show stained cells that have translocated, and also visualizes the pores, which have a nominal diameter of 8 µm. Cells are shown by arrow heads, pores are indicated by arrows. Scale bar: 25 µm. (C) The NC-MT assay was performed with 5% FBS added either to the reservoir, to the insert or into both compartments. After 6 h, cells on the lower membrane surface were fixed, stained and the number of migrated cells was counted. Data were normalized to the maximal migration condition and are shown as means ± SD from three independent experiments. ***: p < 0.001, ns: not significant as determined by one-way ANOVA followed by Dunnett’s post-hoc test (compared to untreated control). (D) Several potential chemoattractants were tested in the NC-MT assay. The concentrations were 5% for fetal bovine serum (FBS), human serum (huSerum), and bovine serum albumin (BSA), 20 ng/ml for fibroblast growth factor 2 and 8 (FGF2/FGF8), glial cell line-derived neurotrophic factor (GDNF), stromal cell-derived factor 1 (SDF-1), and ephrin A1, B1 and B2, 50 µg/ml for basic fibroblast growth factor (bFGF), 10 ng/ml for Wnt3a, 50 ng/ml for platelet-derived growth factor (PDGF), 100 ng/ml for brain-derived neurotrophic factor (BDNF), 1 ng/ml for transforming growth factor β 1 and 3 (TGF-β 1/TGF-β 3), 1 µM retinoic acid (RA), 3 ng/ml for bone morphogenetic protein 4 and 7 (BMP-4/BMP-7), 10 nM for endothelin 1, 100 ng/ml for interleukin-8 (IL-8). UT: untreated. Data are normalized to FBS data and are shown as means ± SD from at least two independent experiments. ***: p < 0.001 as determined by one-way ANOVA followed by Dunnett’s post-hoc test (compared to untreated control). (E) Before use as chemoattractant in the lower (reservoir) chamber (at 5%), FBS was digested with 0.5% pepsin at pH=2.0. The control sample (-pepsin) was run through the same procedure (acidification, 1 h incubation, buffering to pH=7.0), but without pepsin. Activity was determined in the NC-MT assay. Data are normalized to untreated FBS and are shown as means ± SD from three independent experiments. ***: p < 0.001, ns: not significant as determined by one-way ANOVA followed by Dunnett’s post-hoc test (compared to untreated control). (F) FBS samples were heat treated under different conditions, and tested afterwards at 5% concentration in the NC-MT assay for chemotactic activity. Data are normalized to FBS and shown as means ± SD from three independent experiments. ***: p < 0.001, ns: not significant as determined by one-way ANOVA followed by Dunnett’s post-hoc test (compared to untreated control).

The main component of FBS and huSerum, the protein albumin, had no chemotactic activity. However, FBS contains many other proteins and also many small molecules. To get an idea whether a protein is responsible for CTA, we treated FBS in different ways before it was tested for chemoattractant activity. Digestion of proteins by pepsin and protein-denaturation by heating to 70 °C both inactivated the putative chemotaxis-promoting factor (**Figure 1E, F**). From this, we conclude that with a high probability the CTA is at least in part a protein.

### Characterization of the chemotaxis-triggering factor in FBS

As the chemotaxis-triggering factor in FBS is most likely a protein, we wondered whether it could be enriched or even purified. For this purpose, several traditional protein separation strategies were combined in a general strategy. As albumin accounts for >60% of FBS protein, it was important to find a step capable of removing it early on. Different fractional precipitation approaches were tested, and an optimized sequential acetone precipitation was found to be optimal for albumin removal. The second enrichment step was a fast protein liquid chromatography (FPLC) purification with an anion exchange column (HiTrap Q FF). The individual fractions were tested in the NC- MT assay, and those that triggered increased NCC migration were combined, desalted and further purified with another anion exchange column (HiTrap Q HP) (**Figure 2A**). The fractions were tested directly in the NC-MT assay for their migration increasing activity (**Figure S1B**) and we tried to store these for further use. We found that all highly purified fractions lost their CTA bioactivity within 24 h. Multiple approaches of protein stabilization and improved storage were tried. However, we did not identify a procedure that allowed the chemotaxis-promoting factor to be stored overnight once it was highly purified. One potential explanation for this loss of activity is that the “CTA protein” is stabilized by another protein, which is lost upon purification. Due to this situation, the purification and bioactivity testing always had to be performed within one day. As alternative approach to protein chromatography we used ultrafiltration membranes to get an indication on the size range of the CTA contained in FBS. We found that the chemoattractant behaves like a protein with a MW of 50-100 kDa (**Figure 2B**). Mass spectrometric (MS) analysis of the most active fraction purified from the second anion exchange column suggested serpin A1, which has a size of 52 kDa, as potential candidate (**Figure S1C**). Detailed follow-up and confirmation experiments showed that serpin A1 does not have chemotaxis-promoting activity (**Figure S1D**). Analysis of MS spectra showed, that the fraction containing serpin A1, contained at least 20 further proteins (not shown). For this reason, the CTA protein may be easily masked by one of the highly abundant serum proteins, like serpins (Anderson and Anderson, 2002). The stepwise purification, including acetone precipitation and two anion exchangers enabled a 1000- fold purification of the chemotaxis-promoting factor compared to the starting material (**Figure 2C, Figure S1A**). This strong enrichment was not sufficient for MS identification, as even the active fractions contained complex protein mixtures. To solve this problem, additional and more efficient chromatographic columns are necessary. As alternative strategy, we considered a less complex starting material.

**Figure 2:**
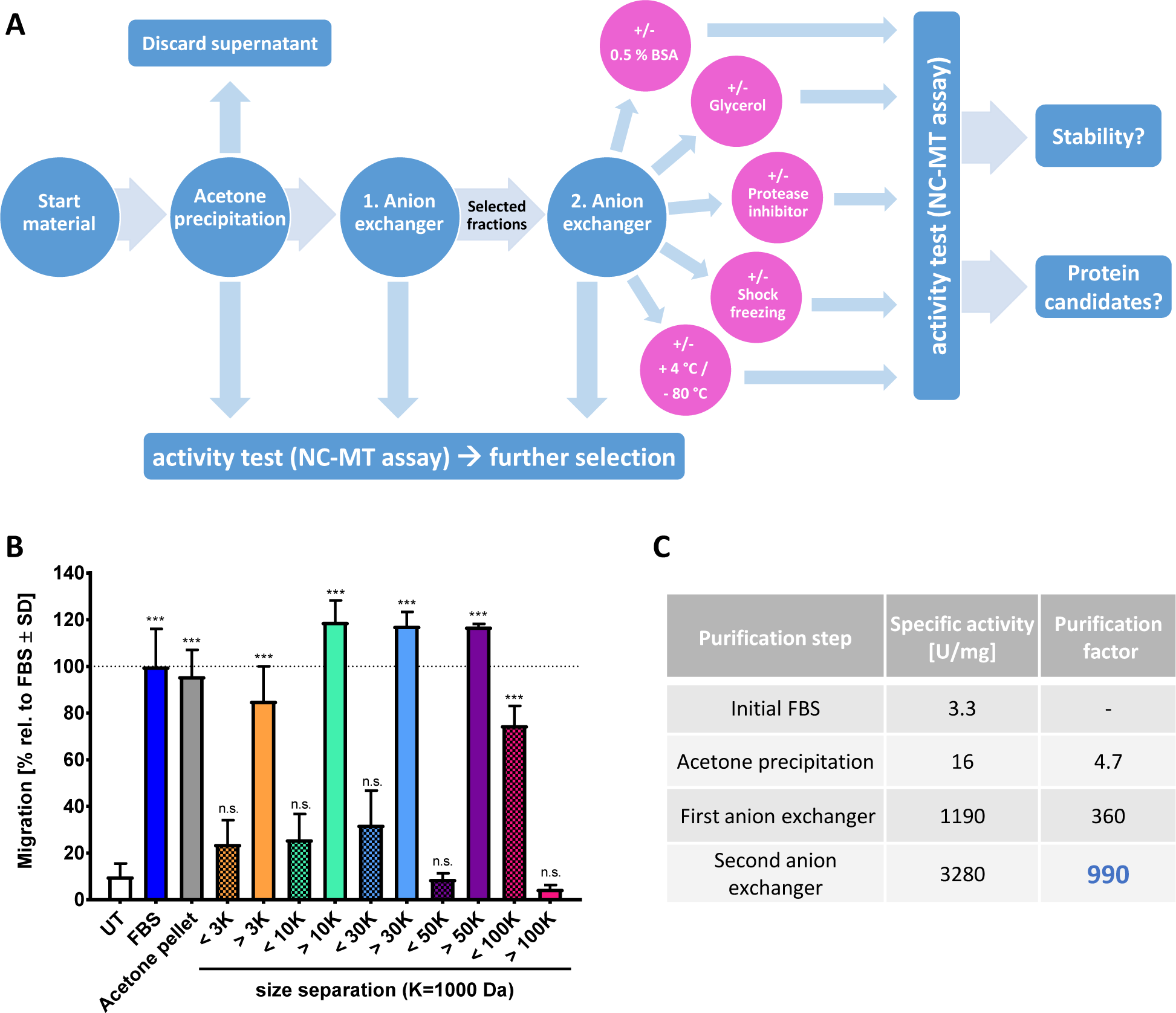
Characterization and purification of the chemotaxis-promoting factor of fetal bovine serum (FBS). (A) Scheme for the enrichment strategy of chemotactic factors from FBS. The activity was measured with the NC-MT assay after each purification step (blue circles). Active fractions from the first anion exchanger (HiTrap Q FF) were combined, desalted and loaded onto the second anion exchange column (HiTrap Q HP). After the last purification step, samples were stored under different conditions (pink circles) for 24 h. Afterwards, the activity of the samples was tested in the NC-MT assay. (B) FBS was precipitated by acetone, and the protein pellet was dissolved and applied to ultrafiltration membranes with different cut-offs. Flow through (<) and filtrate (>) were then tested in the NC-MT- HTS assay. Data are normalized to non-fractionated FBS and shown as means ± SD from three independent experiments. ***: p < 0.001, ns: not significant as determined by one-way ANOVA followed by Dunnett’s post-hoc test (compared to untreated control). (C) The specific activity and the purification factor were determined after each purification step. The used anion exchangers differ in their resin material, loading capacity and flow rates. The most active fractions of each anion exchanger were used for calculations. Data are given relative to the starting material (= FBS).

### Chemotaxis-triggering activity present in conditioned medium of HepG2 cells

We reasoned that all proteins present in serum are produced by cells. Moreover our screen for CTA sources had shown that also human serum is bioactive (**Figure 1D**). Therefore, we set up the hypothesis that some human cells should produce the protein responsible for NCC chemoattractant activity. In order to test this, we used a small cell panel, including HepG2 hepatoma cells, MDA-MB-231 breast adenocarcinoma cells, HeLa cervical cancer cells, HEK-239 human embryonic kidney cells and SH-SY5Y neuroblastoma cells to examine the production of CTA (**Figure 3A**). In a first approach, we used conditioned medium (CM) from all cell lines in the NC-MT assay. The data showed that HepG2 and MDA cells are potent producers of a CTA, HeLa and HEK-239 cells were moderate producers, and SH-SY5Y CM was devoid of any activity. In a second, independent experimental approach, we then confirmed these findings by culturing the cell lines in the lower compartment of the chemotaxis assay setup. SH-SY5Y neuroblastoma cells had no chemoattractant activity at all, i.e. their presence did not trigger any of the NCCs to move through the membrane. This showed that human cells (as such) do not have unspecific chemoattractive effects, if co-cultured in the NC-MT assay. The cells that were chemoattractive for NCCs had a potency order similar to the one found for their CM (**Figure 3A**). Thus, some cells seem to secrete a protein that is chemoattractive for NCCs. We decided to focus on HepG2 as producing cell line. For initial characterization, stability studies were performed on HepG2 CM. Data from these experiments showed that the CTA in this material is completely inactivated by pepsin digestion and by moderate heating (70 °C) (**Figure 3B, C**). These results confirmed that the CTA of HepG2 CM is a protein. Based on this knowledge, a stepwise purification approach was started. Acetone precipitation was used as first step, as it also had a concentrating and de-salting function. Various chromatographic columns were then used. A cation exchanger (HiScreen Capto SP ImpRes) proved to be the most efficient, and it yielded highly enriched CTA (**Figure S2B**). The fractions with the highest migration increasing activity in the NC-MT assay (**Figure S2A**) were used for MS analysis. Fibronectin and apolipoprotein-H were consistently identified in the most active fractions (**Figure S2C, D**). We speculated, that a second chromatographic column would remove one of these two proteins and thus give an indication on which one may trigger migration. Therefore, the active fractions of the cation exchange column were combined, desalted and further purified by an anion exchange column (HiTrap Q HP). The individual fractions were tested in the NC-MT assay and only one fraction triggered NCC migration (**Figure 3D**). Polyacrylamide gel size separation of the CTA containing fraction resulted in two major protein bands (**Figure 3F**) and MS analysis identified them consistently as fibronectin and serum albumin/alpha-fetoprotein (AFP) (**Figure 3G**). As apolipoprotein-H was not present in the active fraction after the second chromatographic purification step, we retained fibronectin as promising candidate and excluded apolipoprotein-H. Serum albumin and AFP share 39% primary structure homology and have the same MW of about 69 kDa (Morinaga et al. 1983). Therefore, MS analysis does not sufficiently distinguish albumin and AFP. However, it was shown that serum deprivation of HepG2 cells increased the production of AFP compared to albumin (Bennett et al. 1998). Moreover, we had found that purified albumin is not chemoattractive. Therefore, we took AFP as another promising CTA candidate. Our assumption was further supported by identification of AFP in HepG2 CM via western blot (data not shown).

**Figure 3:**
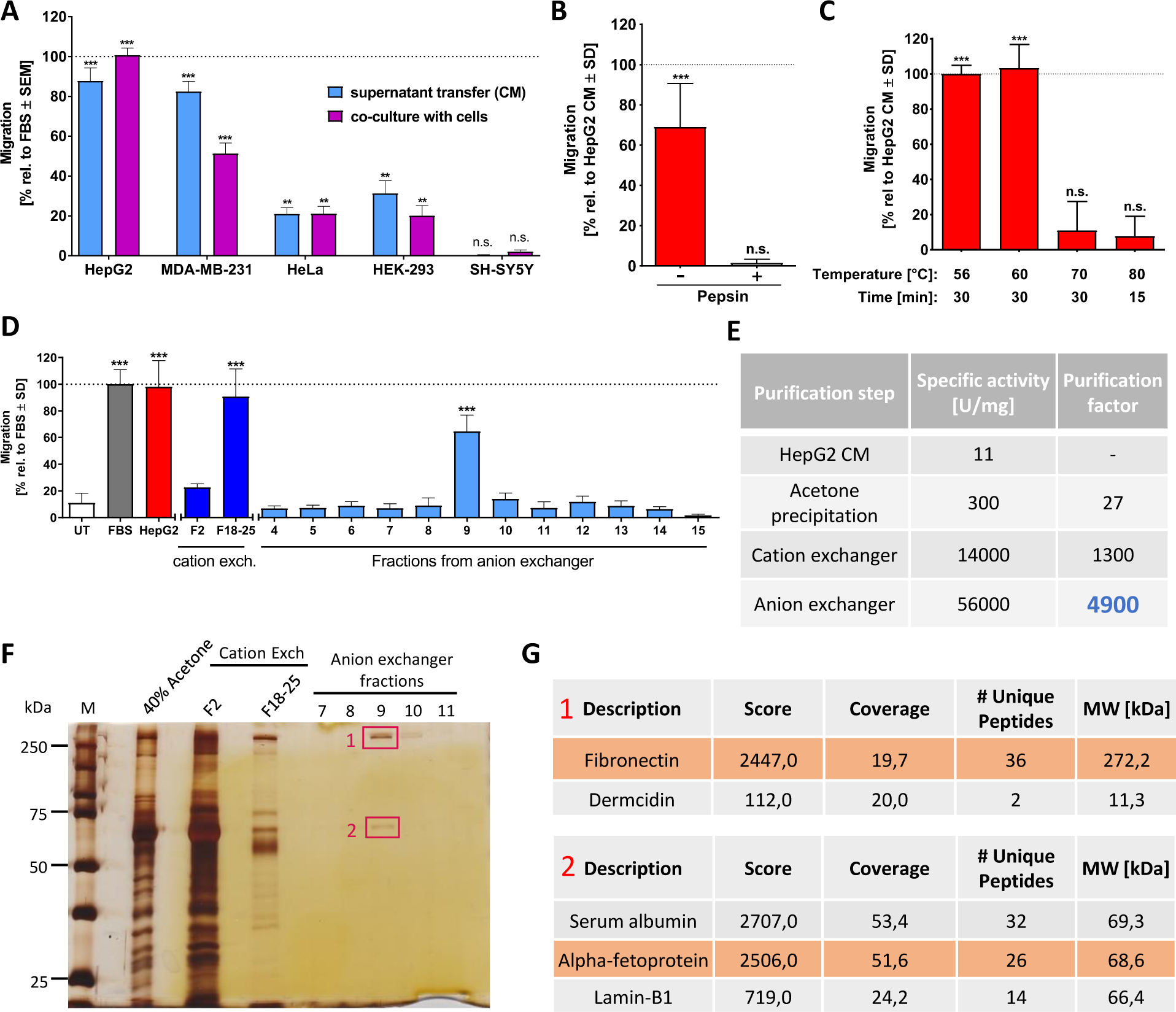
Characterization of the chemotaxis-promoting factor in HepG2 conditioned medium (CM). (A) Production of chemoattractant factors by human cell lines was examined by two experimental set ups: use of conditioned medium (CM) and co-culture. For transfer of CM the indicated cell lines were cultured in medium with 0% FBS for 48 h. The medium supernatant from these cultures was collected, and then used to fill the (lower) reservoirs of the NC-MT assay. The upper insert was filled with NCCs and fresh (unconditioned) medium. For co-culture experiments, cells of the indicated cell lines were grown at the bottom of the reservoir. At the start of the experiment they received fresh medium with 0% FBS, and 48 h later NCCs were added to the transwells above the cell lines in fresh medium (upper compartment). In both cases NCCs were allowed to migrate for 6 h, before the number of cells reaching the lower membrane side was quantified. Data are normalized to the migration triggered by 5% FBS. They are shown as means ± SEM from at least three independent experiments. **: p < 0.01, ***: p < 0.001, ns: not significant as determined by one-way ANOVA followed by Dunnett’s post-hoc test (compared to untreated control). (B) Before use as chemoattractant in the lower (reservoir) chamber, HepG2 CM was digested with pepsin as in Fig. 1E. Data are normalized to HepG2 CM and are shown as means ± SD from three independent experiments. ***: p < 0.001, ns: not significant as determined by one-way ANOVA followed by Dunnett’s post-hoc test (compared to untreated control). (C) HepG2 CM samples were heat treated under different conditions, and tested afterwards at 5% concentration in the NC-MT assay for chemotactic activity. Data are normalized to HepG2 CM and shown as means ± SD from three independent experiments. ***: p < 0.001, ns: not significant as determined by one-way ANOVA followed by Dunnett’s post-hoc test (compared to untreated control). (D) The chemotaxis-triggering factor in HepG2 CM was purified via acetone precipitation followed by a cation exchange column (HiScreen Capto SP ImpRes) and an anion exchange column (HiTrap Q HP). Fraction 2 (F2) and the combined fractions F18-25 (F18-25) from the cation exchanger as well as fractions 4-15 from the anion exchanger were tested at a final concentration of 20% (80% fresh medium) in the NC-MT-HTS assay. As positive control, 5% FBS and 100% HepG2 CM were used. UT: untreated. Data are means ± SD of three independent experiments. *: p < 0.05, **: p < 0.01, ***: p < 0.001 as determined by one-way ANOVA followed by Dunnett’s post-hoc test (compared to untreated control). (E) The specific activity and the purification factor were determined after each purification step. (F) Samples from acetone precipitation (40% acetone), fraction 2 (F2) and combined fractions 18-25 (F18-25) from the cation exchanger as well as fractions 7-11 from the anion exchanger were separated on a 10% SDS gel, and bands were visualized by silver staining. Bands cut out for MS analysis are marked in red. 1: Fibronectin, 2: Alpha-fetoprotein. (G) Proteins in fraction 9 from the anion exchanger were separated on a 10% SDS gel and two bands were cut out for mass spectrometry (MS) analysis. The table represents the MS result of bands 1 and 2 of fraction 9, detecting fibronectin and alpha-fetoprotein as the most abundant proteins. MS analysis was performed from two independent experiments with the same results.

For further narrowing down the identity of the CTA, we decided to focus on the two most abundant proteins present in the active fraction, fibronectin and AFP. To probe the role of fibronectin, it was removed from HepG2 CM by an affinity precipitation, using gelatin sepharose beads (**Figure S3C, D**). Testing in the NC-MT assay showed that HepG2 CM without fibronectin has the same migration increasing effect on NCCs as HepG2 CM with fibronectin (**Figure S3B**). We therefore excluded fibronectin as the potential chemoattractant factor in HepG2 CM. Two further proteins identified by MS were excluded as likely contaminants: Dermcidin is present in human sweat and is therefore often found in MS samples (**Figure 3G**). Also, the nuclear lamina protein lamin-B1 was discarded as likely candidate (**Figure 3G**). As SH-SY5Y neuroblastoma cells had no chemoattractant activity at all in the NC-MT assay (**Figure 3A, Figure S4A**), and CM produced from these cells did not contain albumin or AFP (**Figure S4B**), we expressed recombinant AFP in the neuroblastoma cell line. Engineered SH-SY5Y cells produced AFP (**Figure S4D,E**), but did not show chemotactic activity in the NC-MT assay (**Figure S4C**). We concluded form these data, that AFP is not the chemotaxis-promoting factor in HepG2 CM. Thus, our purification approach, which resulted in a 5000-fold enrichment of the starting material (**Figure 3E**), did not allow the CTA identification. However, we were able to enrich a definitely human NCC chemotaxis factor to a high degree. Active HepG2 fractions contained clearly less protein than active FBS fractions, but the supernatant production was very resource-requiring. Moreover, it cannot be excluded that cancer cells produce a factor that is not physiologically relevant. Therefore, we considered other sources.

### Human platelet lysate as animal-free CTA alternative

Human platelet lysate is a high quality human-derived product known to be rich in growth factors. It appeared as optimal alternative starting material, as it is commercially available. In a pilot experiment, we produced a small amount of huPL ourselves and observed potent bioactivity in the NC-MT assay (not shown). To follow up on this, we obtained huPL from various suppliers. We were surprised to observe that the huPL contained large amounts of albumin (**Figure S5A**). Our investigations showed, that plasma is added to all commercial huPLs. The suppliers argued that this is necessary to stabilize the platelet factors and to guaranty optimal cell growth when huPL is used as cell culture additive (Chou and Burnouf 2017; Horn et al. 2010). When we tested different lots of commercially available huPLs in the NC-MT assay, we found that this material potently triggered NC migration (**Figure 4A**). In a next step, we used our established procedures to verify that a protein of the lysate is responsible for chemotaxis. Data from these experiments showed that the CTA in huPL is inactivated by pepsin digestion and by heating (**Figure 4B, C**). We therefore concluded that the chemotaxis-promoting factor in huPL is a protein, similar to the factor present in FBS and HepG2 CM. For purification of the CTA, we optimized the strategy and started with a chromatographic purification step using the HiScreen Capto SP ImpRes cation exchange column (**Figure 4D, Figure S5B**). The acetone precipitation was performed afterwards on the pooled active fractions obtained. This way, the pellet could be frozen and stored at -20 °C. Through this improved procedure, it was no longer necessary to perform all purification steps within one day. Additionally, many acetone pellets could be combined and a large batch could be produced for further purification. A disadvantage of this process was that the pellet was hard to dissolve, and some material was lost. By running the re-dissolved material over an anion exchange column (HiTrap Q HP), the CTA could eventually be purified up to 2000-fold, compared to the starting material (**Figure 4E**).

**Figure 4:**
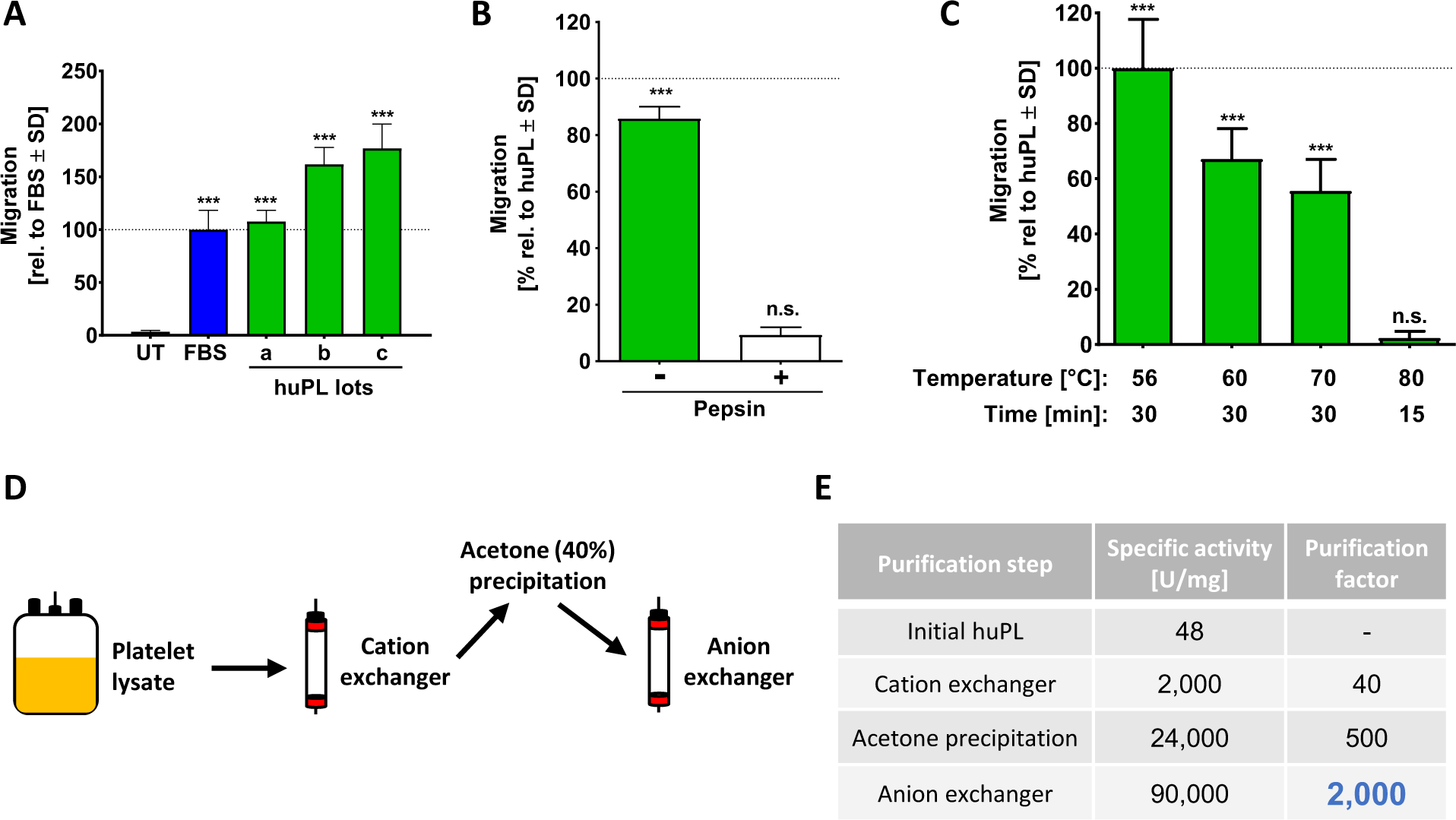
Characterization of the chemotaxis-promoting factor from human platelets. (A) Several commercially available huPLs (lots a-c) were tested at a final concentration of 5% in the NC-MT assay. Heparin (at a final concentration of 2 U/ml) was added to lots b and c to prevent coagulation of the medium. FBS (5%) was used as positive control. Data are normalized to FBS and are shown as means ± SD from three independent experiments. ***: p < 0.001 as determined by one-way ANOVA followed by Dunnett’s post-hoc test (compared to untreated control). (B) Before use as chemoattractant in the lower (reservoir) chamber (at 5%), huPL was digested with 0.5% pepsin (as in Fig. 1E). Data are normalized to huPL and are shown as means ± SD from three independent experiments. ***: p < 0.001, ns: not significant as determined by one-way ANOVA followed by Dunnett’s post-hoc test (compared to untreated control). (C) huPL samples were heat treated under different conditions, and tested afterwards at 5% concentration in the NC-MT assay for chemotactic activity. Data are normalized to huPL and shown as means ± SD from three independent experiments. ***: p < 0.001, ns: not significant as determined by one-way ANOVA followed by Dunnett’s post-hoc test (compared to untreated control). (D) Scheme for the enrichment strategy for chemotactic factors from human platelet lysate (huPL). Platelet lysate was fractionated on a cation exchange column (HiScreen Capto SP ImpRes), followed by an acetone precipitation and analysis of the pellet material on an anion exchange column (HiTrap Q HP). (E) The specific activity and the purification factor were determined after each purification step.

### Time-lapse analysis of the chemotactic activity of FBS, HepG2 CM and huPL

Having established three ways to obtain a potent NCC chemotactic factor in a population-based assay (NC-MT), we were interested to obtain additional and more direct proof of chemotactic activity on individual cells. The NC-MT assay determines the number of migrated cells at the end of the assay and cells are not observable during the migration. To avoid these issues we performed the µ-slide chemotaxis assay. This setup uses a complex cell culture format (produced by ibidi), in which cells can be placed in a stable chemical gradient, and observed over several hours. (**Figure 5A**, **Figure S6A**). The cells were observed via time-lapse imaging, and the migration tracks of individual cells were visualized. Initial controls showed that NCC migrated in all directions when there was no chemoattractant present (**Figure 5B left**). When cells were exposed to the same concentration of a chemoattractant on both sides, they migrated towards both stimuli with about the same frequency (**Figure 5B right**). For classical chemotaxis testing, only one reservoir was filled with a chemoattractant (FBS, HepG2 CM or huPL) so that cells were within a stable gradient. Under these conditions, they migrated towards the higher concentration of the chemoattractant (**Figure 5C**).

**Figure 5:**
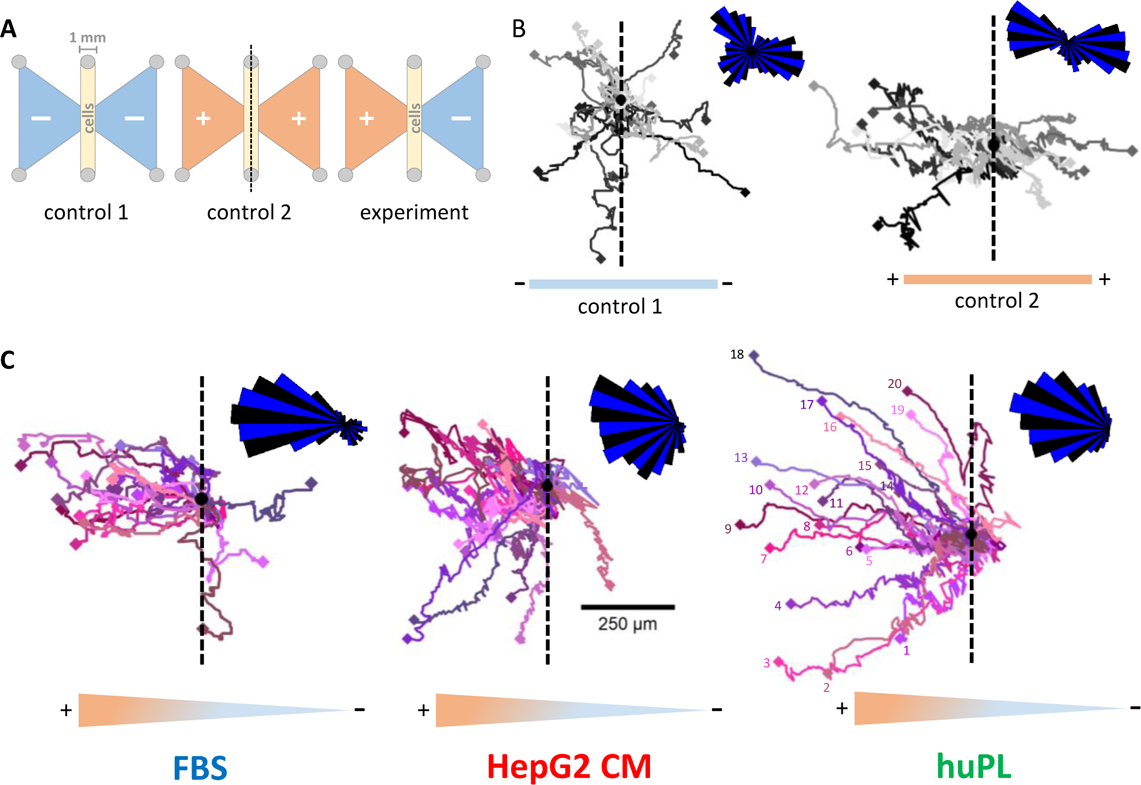
Tracking of single cells to compare chemotactic activity of FBS, huPL and HepG2 CM. (A) Graphical illustration of the µ-slide chemotaxis assay setup, as seen from above (ibidi). Cells were seeded in the 1 mm broad gap between the reservoirs (oblong middle compartment). Reservoirs on both sides (shown as triangles) were either filled with medium (-) or chemoattractant dissolved in medium (+). (B) Representative cell trajectories (recorded for 24 h) of 20 NCCs with medium in both reservoirs (control 1, left) or FBS (5%) in both reservoirs (control 2, right). For quantification of the migration, the chamber was placed onto a microscope stage, and cell movement was observed by time-lapse video imaging. The black dot is the starting point of all cells in a re-normalized co-ordinate system (overlay of all cell starting points). The dashed line was inserted as visual aid to symbolize the “watershed” of the gradient. The density plot in the upper right corner was constructed by separating the migration area into 36 sectors (each 10° wide) starting from the migration origin. The length of the segments indicates the distribution of cell counts inside the sectors. For visualization reasons, sectors were alternatingly stained blue and black. (C) Representative cell trajectories (recorded for 24 h) of 20 NCCs with medium in the right reservoir and FBS, huPL or HepG2 CM filled in the left reservoir. Numbers for each individual cell are only indicated in the right figure for graphical reasons.

The software provided by the assay chamber supplier allowed quantification of migration parallel or perpendicular to the gradient (**Figure S6A**). Chemotaxis is defined by this program as a form of migration that is more effective parallel to the gradient than perpendicular to it. Addition of FBS, HepG2 CM or huPL to one reservoir, led to clear chemotaxis (**Figure S6B**). The results confirmed that FBS, HepG2 CM and huPL trigger chemotaxis in individual NCCs. Moreover, NCCs in a HepG2 CM or huPL gradient migrated also longer distances and faster, compared to the control cells (**Figure S6D**). In conclusion, FBS, HepG2 CM and huPL are verified sources of CTA for NCCs.

## Conclusion and Outlook

Altogether, we have shown here that it is possible to establish an *in vitro* assay for directed migration of human neural crest cells (chemotaxis), a pivotal process in fetal development. Human material can be exclusively used for the assay, and it goes beyond the assessment of non-directed movement capacity (chemokinesis). Thus, it can be used to study processes and factors (pathological, toxicological or pharmacological) that specifically affect the sensing of a chemoattractive gradient and accordingly the directed movement of NCCs. We also demonstrated, that the assay allows a highly quantitative readout, as it was successfully used here for determining the bioactivity spectrum of chromatographic fractions or of CTA preparations after various treatments (heat, proteases, etc.). This is a key feature, important for the development of new approach methods (NAMs) that can be used for testing of potential developmental toxicants (in particular developmental neurotoxicants). In the past, tremendous research effort has been invested in animal models of neural crest migration, and the search for human-relevant chemotactic factors has been neglected. For this purpose, ground work, as described here, is essential for providing a basis of a new generation of tests, based only on human cells and human relevant material and processes (here chemoattractant factors, cell culture coating, cell culture medium). Even though we have not yet succeeded in identifying a protein that can be produced recombinantly, and further used for such assays, we have provided here protocols, which show how suitable protein fractions can be produced to establish a NAM.

We described here three sources of CTA: HepG2 supernatant, huPL, and serum. For the latter source, most work focused on FBS, but we showed that also human serum is bioactive. Indeed, we do not have a perfect proof that the chemotaxis-promoting factor in huPL is a platelet protein. Commercially-available huPL always contained serum proteins. As an alternative approach, we produced purified platelets ourselves. These contained distinctly less plasma proteins than commercial huPL, but we cannot completely exclude contaminations.

Possibly, there is not only one protein with CTA, and such factors may work independently, or they may work synergistically. We believe that there is strong evidence for at least one factor present in serum. First, because we find CTA in cell-free serum; second, because HepG2 cells, which are known to produce serum proteins (Franko et al. 2019; Trefts et al. 2017) produce such a factor. Whether the factor found in serum and the one in HepG2 supernatants (or huPL) is the same protein cannot be decided on the basis of the available data.

For better defining the protein factor, additional efforts are necessary. The combination of better chromatographic approaches together with extensive mass spectrometric characterization of all fractions may provide a step forward. However, this strategy can only be fully developed, if it is possible to stabilize, and store highly bioactive fractions. At present, the loss of bioactivity of the most active fractions within 24 h makes purification strategies extremely challenging. Additionally, increasing the amount of starting material is limited by the binding capacity of the chromatographic columns. To circumvent these limitations, removal of unwanted proteins from the starting material and simultaneous concentration of the remaining proteins (e.g. by a precipitation step) is necessary. Another issue is the potentially high bioactivity of a chemotactic factor together with its extreme dilution by other proteins. If the factor has hormone/cytokine-like properties, it is likely to have low nM or even pM affinities and may therefore only be present in pM concentrations. Such a protein might be easily masked by highly abundant serum proteins, e.g. upon gel separation. Such issues are well-known in biology and some of the most obvious and important bioactive molecules could never be purified by traditional methods. This applies e.g. to the erythropoietin receptor, the corticotrophic hormone receptor or the tumor necrosis factor receptor, which were all eventually identified by functional expression cloning (Beutler et al. 1985; Chen et al. 1993; D’Andrea et al. 1989). Such strategies may be used in the future for identification of chemotaxis factors, considering that the bioassay works at relatively high throughput.

The above purification strategies are of mid-term and long-term interest, as they inform on the underlying human physiology. They may also offer some advantage for assay development and application. However, a purified factor is by no means necessary to go ahead. Biology has a long and successful tradition of using complex materials, often not fully defined in their composition, for quantitative assays. We have shown here procedures to purify the CTA several thousand-fold, which is already an intermediate step towards a more defined material. Most importantly, we demonstrated how FBS could be exchanged for fully human material, and on this basis, a complete humanized NAM can be established.

## Acknowledgements

This work was supported by CEFIC, the BMBF, EFSA, and the DK-EPA (MST-667-00205). It has received funding from the Land-BW (NAM-ACCEPT) and the European Union’s Horizon 2020 research and innovation program under grant agreements No. 681002 (EU-ToxRisk), No 964537 (RISK-HUNT3R), No. 964518 (ToxFree) and No. 825759 (ENDpoiNTs).

The Proteomics Centre of the University of Konstanz is acknowledged for providing excellent support.

## Abbreviations

cMINC: circular
MINC DMSO: dimethyl sulfoxide
DNT: developmental neurotoxicity
ECM: extracellular matrix
EGF: epidermal growth factor
FBS: fetal bovine serum
FGF: fibroblast growth factor
FPLC: fast protein liquid chromatography
HTS: high-throughput screening
huPL: human platelet lysate
iPSC: induced pluripotent stem cell
MINC: migration inhibition of neural crest cell
MW: molecular weight
NAM: new approach method
NCC: neural crest cell
NC-MT: neural crest membrane translocation
NGRA: next generation risk assessment
PBS: phosphate buffered saline
PCB: polychlorinated biphenyl
PDGF: platelet derived growth factor
RA: retinoic acid
SDF-1: stromal cell derived factor 1
VEGF: vascular endothelium growth factor
VPA: valproic acid

## Supplementary information

**Figure S1:**
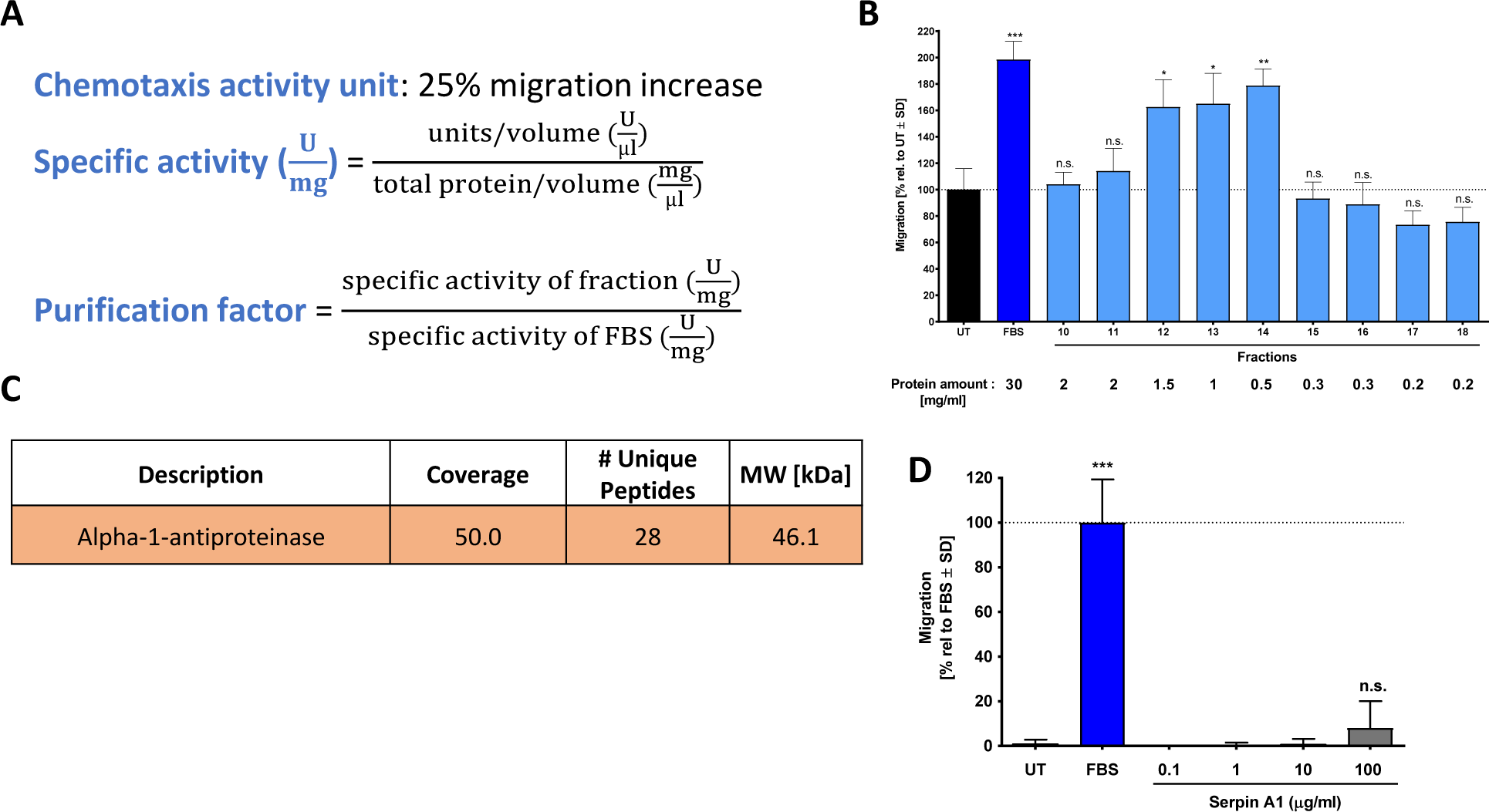
Characterization and purification of the chemotaxis-promoting factor of bovine serum and exclusion of serpin A1 as chemoattractant candidate. (A) Definition and calculation of chemotaxis activity unit, specific activity and purification factor. One unit is defined as the migration increase of NCCs in the cMINC assay within 24 h migration time, as well as in the NC- MT assay within 6 h migration time. (B) FBS was purified via acetone precipitation and two anion exchangers. Fractions of the second anion exchanger (HiTrap Q HP) were tested at a final concentration of 5% in the cMINC assay for their migration increasing activity. As positive control 5% FBS was used. UT: untreated. Data are normalized to untreated control and shown as means ± SD of two independent experiments. *: p < 0.05, **: p < 0.01, ***: p < 0.001 as determined by one-way ANOVA followed by Dunnett’s post-hoc test (compared to untreated control). (C) Fraction 14 of the second anion exchanger (HiTrap Q HP) indicated the highest migration increase. The fraction was separated on a 10% SDS gel and analysed by mass spectrometry (MS) analysis. The table represents the MS result of fraction 14 detecting serpin A1 as the most abundant protein. (D) The most abundant protein serpin A1 present in the fraction with the highest migration activity, was tested at the indicated concentrations in the NC-MT assay. Data are normalized to FBS and shown as means of five technical replicates ± SD. ***: p < 0.001 as determined by one-way ANOVA followed by Dunnett’s post-hoc test (compared to untreated control).

**Figure S2:**
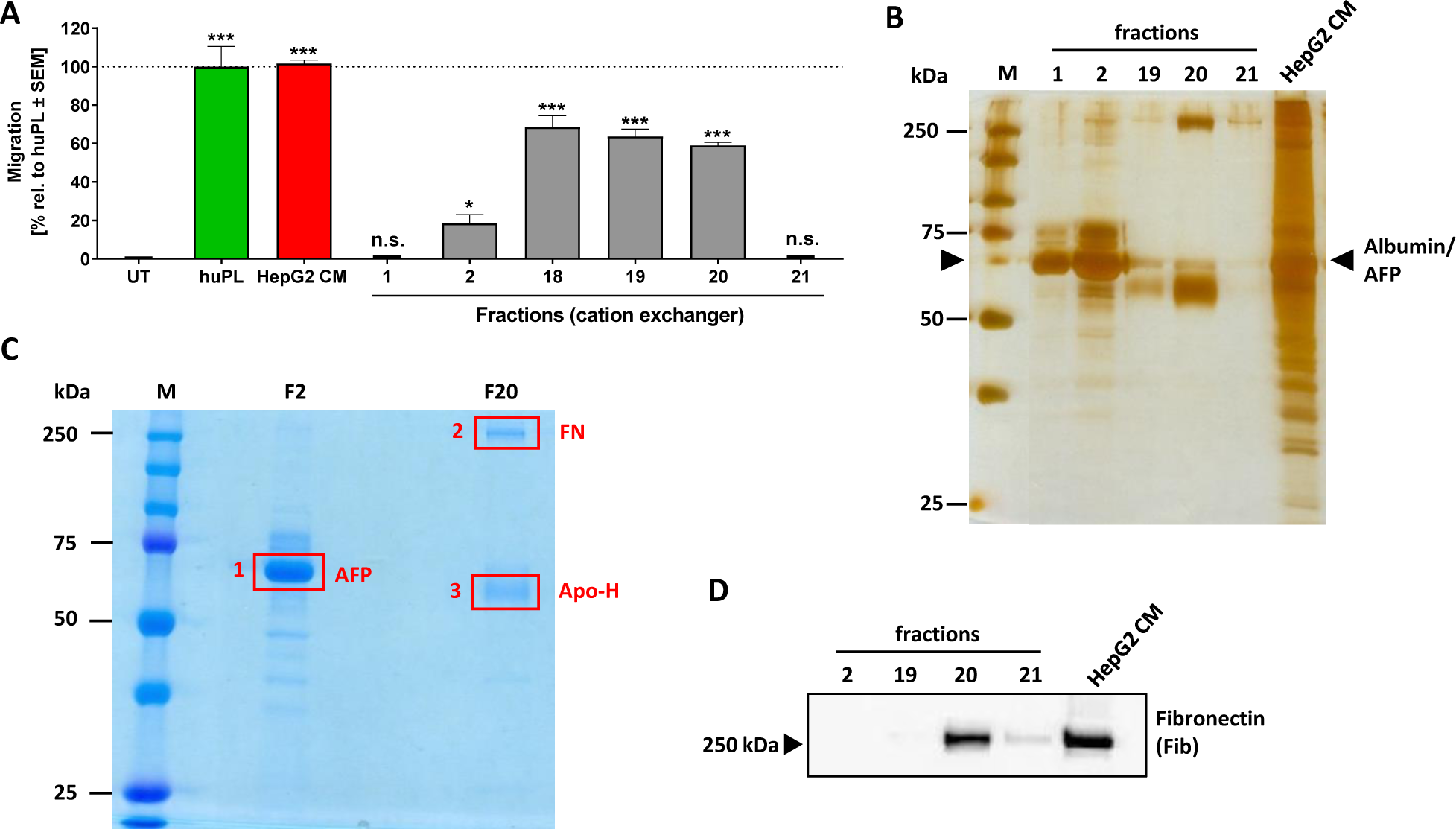
Characterization of chemotaxis-promoting factor in HepG2 CM after acetone precipitation and cation exchanger. (A) HepG2 CM was precipitated by acetone, followed by a cation exchange column (HiScreen Capto SP ImpRes). Fractions 1 and 2 (flow-through) and fractions 18-21 of the cation exchanger were tested at a final concentration of 20% in the NC-MT-HTS assay. As positive control 5% huPL and 100% HepG2 CM were used. UT: untreated. Data are normalized to 100% HepG2 CM and are shown as means ± SEM of two independent experiments *: p <0.05, ***: p < 0.001 as determined by one-way ANOVA followed by Dunnett’s post-hoc test (compared to untreated control). (B) Samples from HepG2 CM and fractions 1, 2, and 19-21 of the cation exchange column were separated on a 10% SDS gel, and bands were visualized by silver staining. (C) Samples of fraction 2 and fraction 20 of the cation exchanger were separated on a 10% SDS gel, and bands were visualized by coomassie staining. Bands cut out for MS analysis are marked in red. The most abundant proteins, detected in each band by MS analysis are stated. 1: Alpha-fetoprotein (AFP); 2: Fibronectin (FN); 3: Apolipoprotein-H (Apo-H). (D) Samples from HepG2 CM and fractions 2, 19, 20 and 21 of the cation exchange column (HiScreen Capto SP ImpRes) were separated on a 10% SDS gel and afterwards transferred onto a nitrocellulose membrane. Western blot analysis using anti-fibronectin antibody was performed.

**Figure S3:**
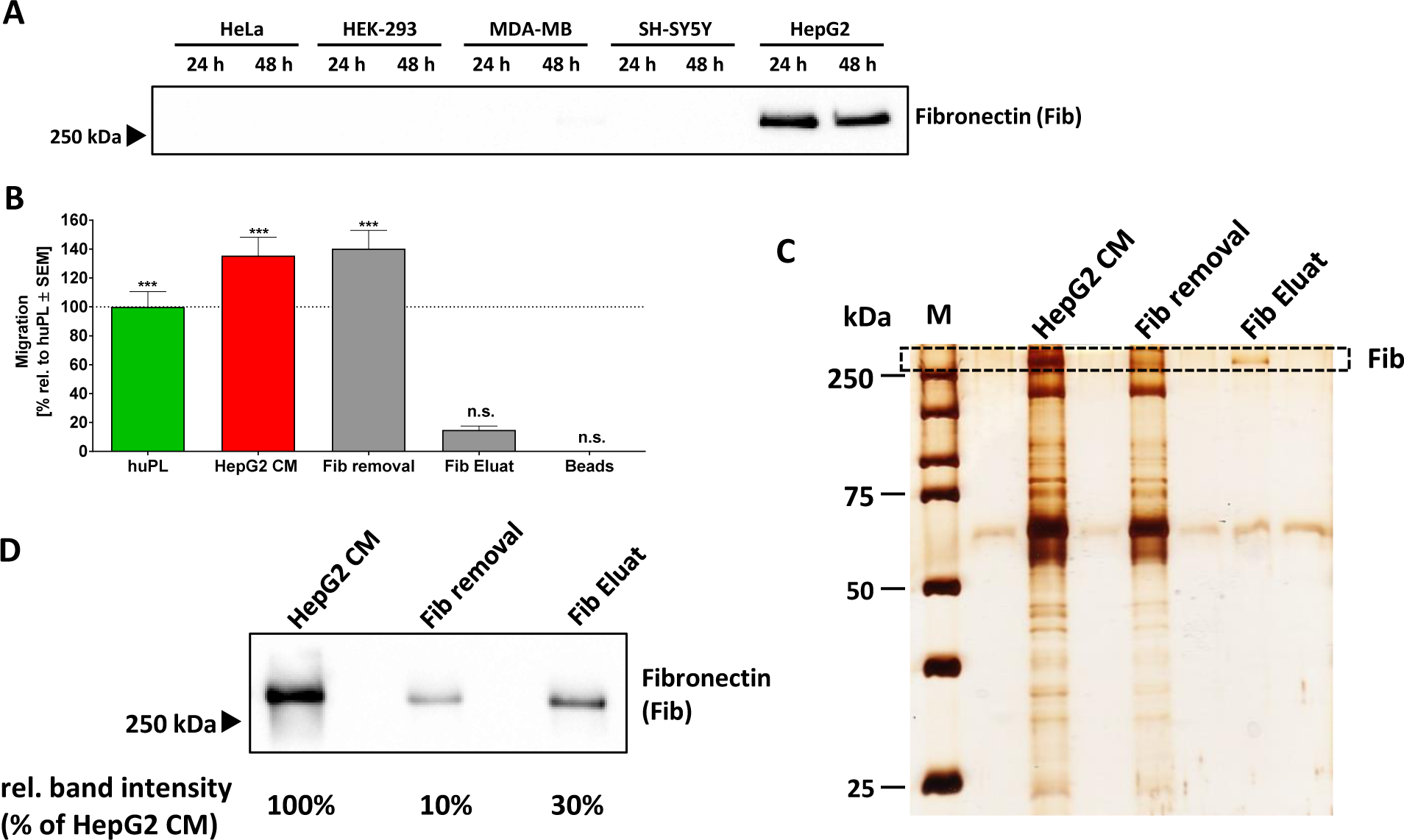
Exclusion of fibronectin as chemoattractant factor in HepG2 conditioned medium. (A) Conditioned medium was produced from the indicated cell lines. Samples were taken after 24 h and 48 h of starvation, separated on a 10% SDS gel, and afterwards transferred onto a nitrocellulose membrane. Western blot analysis using anti-fibronectin antibody was performed. (B) The Gelatin Sepharose 4B beads (Cytiva) were mixed with 5 mg/ml BSA to block nonspecific protein binding, centrifuged at 100 x g for 3 min and afterwards the supernatant was discarded. This procedure was repeated with 6 M urea solution followed by two washing steps with PBS, all at room temperature. The washed gelatin beads were mixed with 10 ml of HepG2 CM and incubated for 24 h at 4 °C on a tube rotator. The next day the gelatin beads with the bound fibronectin were centrifuged at 100 x g for 3 min and the supernatant was transferred into a fresh tube. New gelatin beads were added to the HepG2 CM and incubated for another 24 h at 4 °C on a tube rotator. After centrifugation, the HepG2 CM without fibronectin (Fib removal) was transferred to a new tube and tested at a concentration of 100% in the NC-MT assay. The gelatin beads were washed with 6 M urea solution to remove the bound fibronectin and centrifuged at 100 x g for 3 min. The supernatant containing fibronectin (Fib Eluat) was centrifuged through a 10 kDa cut-off filter, and washed once with PBS to remove the urea solution, before it was tested at a final concentration of 20% in the NC-MT assay. Data are normalized to 5% huPL and shown as means ± SEM from three independent experiments. ***: p < 0.001 as determined by one-way ANOVA followed by Dunnett’s post-hoc test (compared to untreated control). (C) Samples from HepG2 CM before (HepG2 CM) and after fibronectin removal (Fib removal) as well as fibronectin eluated from the beads (Fib Eluat) were separated on a 10% SDS gel and bands were visualized by silver staining. (D) Samples from HepG2 CM before (HepG2 CM) and after fibronectin removal (Fib removal) as well as fibronectin eluated from the beads (Fib Eluat) were separated on a 10% SDS gel and afterwards transferred onto a nitrocellulose membrane. Western blot analysis using anti-fibronectin antibody was performed.

**Figure S4:**
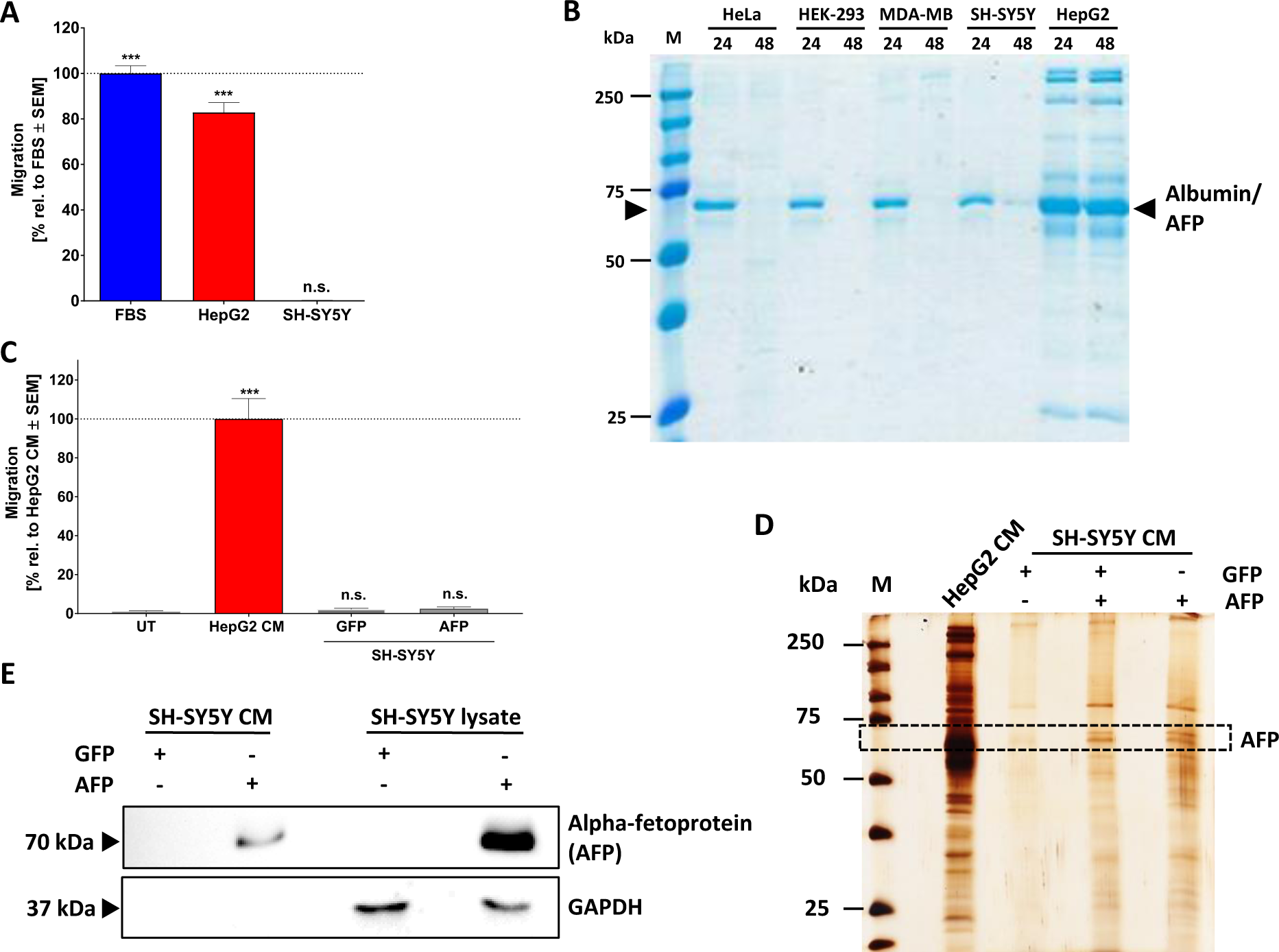
Exclusion of alpha-fetoprotein (AFP) as chemoattractant factor in HepG2 conditioned medium. (A) HepG2 and SH-SY5Y cells were starved with medium containing 0% FBS for 48 h. The conditioned medium was added at a concentration of 100% to the reservoirs of the NC-MT assay and NCCs were allowed to migrate for 6 h. Data are normalized to 5% FBS and shown as means ± SEM from three independent experiments. ***: p < 0.001 as determined by one-way ANOVA followed by Dunnett’s post-hoc test (compared to untreated control). (B) Conditioned medium was produced from the indicated cell lines. Samples were taken after 24 h and 48 h of starvation, separated on a 10% SDS gel, and bands were visualized by coomassie staining. (C) Cells were transfected with Lipofectamine™ 3000 (Thermo Fisher Scientific). Therefore, cells were seeded in 6-well plates and grown until they reached 70% confluency. For transfection, 5 µg DNA was mixed with Opti-MEM™ medium, Lipofectamine™ 3000 reagent, P3000™ reagent, and incubated for 15 min at room temperature, before the DNA-lipid complex was added to the cells. After 24 h of incubation, the medium was aspirated, fresh medium with 0% FBS was added, and the cells were incubated for another 24 h. The conditioned medium was harvested and centrifuged at 314 x g for 4 min, to remove cell debris. The alpha 1 fetoprotein clone IRATp970C1243D was obtained from Source BioScience Cambridge. Conditioned medium was tested in the NC-MT assay at a final concentration of 100%. UT: untreated. Data are normalized to 100% HepG2 CM and shown as means ± SEM from three independent experiments. ***: p < 0.001 as determined by one-way ANOVA followed by Dunnett’s post-hoc test (compared to untreated control). (D) Samples from SH-SY5Y CM of cells transfected with AFP, GFP or both were separated on a 10% SDS gel, and bands were visualized by silver staining. (E) Samples from SH-SY5Y CM and lysate of cells transfected with either AFP or GFP as control were separated on a 10% SDS gel, and afterwards transferred onto a nitrocellulose membrane. Western blot analysis using anti-AFP and anti-GAPDH antibody was performed.

**Figure S5:**
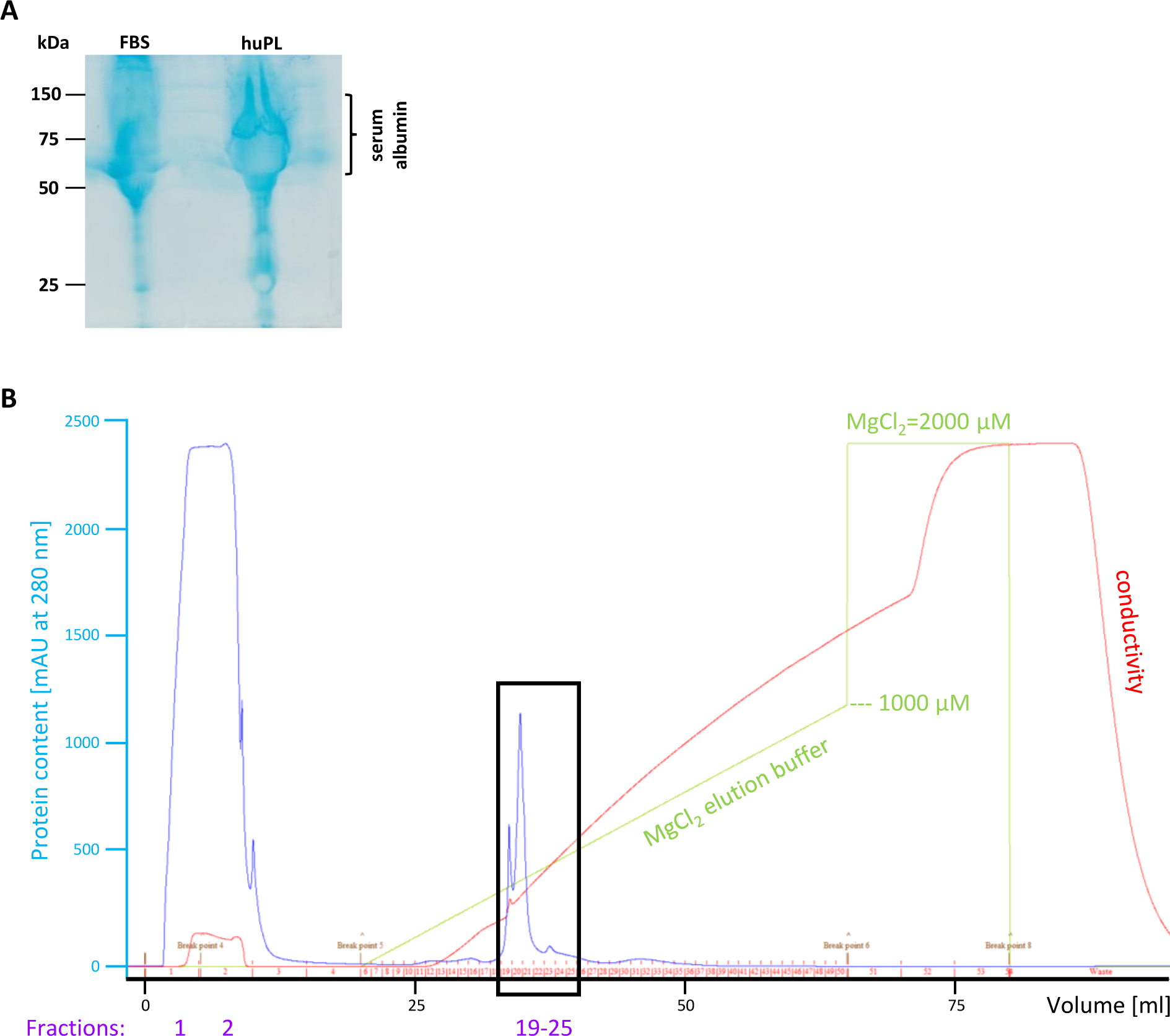
Characterization and purification of the chemotaxis-promoting factor in huPL. (A) FBS and commercial huPL (both 30 mg/ml) were fractionated by polyacrylamide gel electrophoresis (PAGE). Each lane was loaded with 20 µl of sample. For visualization of the protein, gels were stained with coomassie. MW markers were run in parallel (lane not shown, but marker positions are indicated to the left). The large content of serum albumin, indicated by a smear at MW > 55 kDa, prevented separation and visualization of proteins with MW > 50 kDa. (B) Typical elution profile of huPL on a cation exchange column (HiScreen Capto SP ImpRes). The protein content is indicated in blue, the conductivity is shown in red and the concentration of the elution buffer (MgCl_2_) is depicted in green. The numbers of the collected fractions are given at the bottom in violet. The column was loaded with 5 ml huPL and then washed with 20 ml (5 x the volume of the column) 10 mM Tris pH 7.4. All unspecifically bound proteins were washed off, shown in the first peak (fraction 1/2) together with a small salt peak. The elution started with a gradient of 1-1000 mM MgCl_2_, followed by a high salt washing step with 2 M MgCl_2_. Fractions 19-25 triggered chemotaxis in the NC-MT assay (black rectangle).

**Figure S6:**
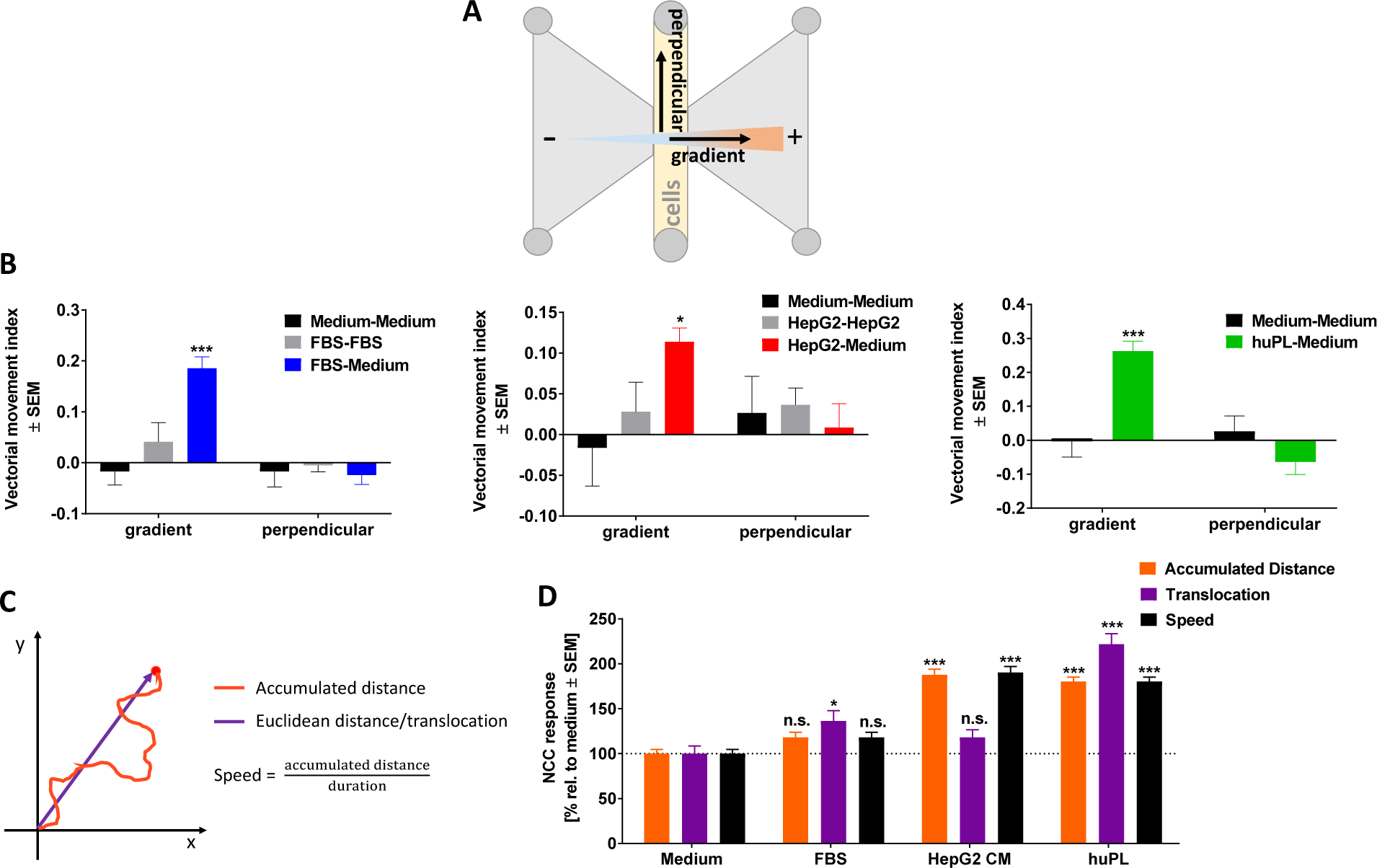
Single cell migration behaviour in µ-slide chemotaxis chambers. (A) Graphical representation of the vectorial movement index. This index can be parallel to the gradient (gradient) or perpendicular to the gradient (perpendicular). A chemotactic effect is fulfilled if gradient > perpendicular for the chemoattractant-medium (+/-) condition, whereas for the control conditions medium/medium (-/-) and chemoattractant/chemoattractant (+/+) both vectorial movement indices are around 0. (B) Calculation of vectorial movement index gradient and vectorial movement index perpendicular for FBS, huPL and HepG2 CM treated cells. Data are shown as means ± SEM of at least two independent experiments *: p <0.05, ***: p < 0.001 as determined by two-way ANOVA followed by Sidak’s post-hoc test (compared to Medium-Medium conditions). (C) Graphical representation and definition of accumulated distance, translocation (euclidean distance from start to endpoint of track) and speed (distance along the track per time). (D) Cells were treated with medium, FBS, huPL and HepG2 CM and time-lapse imaging was performed for 24 h. Accumulated distance, translocation and speed were calculated from the trajectory of 20 cells from two independent experiments. *: p <0.05, ***: p < 0.001 as determined by one-way ANOVA followed by Dunnett’s post-hoc test (compared to medium; performed separately for each endpoint).

